# Genetic Programming of Bacterial Microcompartments: Operon Order as a Tool for Nanoscale Morphogenesis

**DOI:** 10.64898/2026.05.01.722140

**Authors:** Dimple Goel, Preeti Negi, Aarcha Radhakrishnan, Sharmistha Sinha

## Abstract

Gene order is a powerful design principle for protein nanomachines. In nature, gene organisation ensures the precise assembly of functional protein nanostructures. We demonstrate how genetic repositioning of the key structural gene *pduN*, within the operon encoding a self-assembling protein nanocompartment, sculpts the morphology and function of bacterial microcompartments (BMCs). Relocating *pduN* to new operonic positions dramatically altered the size, shape, and catalytic output of BMCs, despite identical protein sequences. These shifts reveal how gene order may control nanoscale assembly and compartmentalised function. Our findings establish operon architecture as a programmable genetic framework for nanostructure morphogenesis and provide a synthetic biology strategy to engineer self-assembling nanodevices with customised geometries and activities.

## Introduction

In biological systems, the assembly of functional nanostructures is guided by genetic organization, where gene order serves as an important regulatory feature. Across diverse lineages, the physical organization of genes within operons has evolved as a means of co-regulation and as a fine-tuned system to ensure the correct stoichiometry, timing, and spatial assembly of complex molecular nanomachines^1–7^. Yet, despite its evolutionary importance, the role of gene order as a programmable tool for controlling nanoscale architecture remains unexplored. Bacterial operons encoding self-assembling protein nanocompartments, bacterial microcompartments (BMCs), offer an ideal model to probe this connection. These proteinaceous nanocages encapsulate enzymes within selectively permeable shells, enabling spatially confined biochemical reactions ^8–11^. One striking example is the BMCs, polyhedral proteinaceous assemblies of size 80-100 nm that encapsulate enzymes within a selectively permeable shell, coordinating metabolic flux with molecular containment^12^ **(Figure 1ai and aii)**. Among these, the 1,2-propanediol utilization (PduBMC) exemplifies genetically compact nanostructure design, with >20 genes forming a contiguous operon that directs autonomous assembly into polyhedral microcompartments^13^. Key genes encode for the shell proteins (e.g *pduA, pduB, pduB’, pduM, pduN, pduJ, pduK, pduT*, and *pduU*) that make the outer shell and the enzymes (e.g., *pduCDE, pduL, pduP, pduQ*, and *pduW*) that metabolize 1,2-propanediol (1,2 PD) while sequestering toxic intermediates ^14^ **(Figure 1bi and Sup Figure 1a)**. The *pdu* operon can be simplified based on shell protein encoding gene clusters (SPC) and enzyme encoding gene clusters (EC), where the enzyme genes can be seen interspaced between shell protein genes **(Figure 1cii)**. Further, similar operonic positioning of key structural genes is observed across multiple BMC systems, consistent with a potential functional role^15^. For instance, *pduN*, encoding the vertex protein that caps BMC polyhedra^16^, resides mid-operon, unlike upstream-positioned major shell genes^13^ **(Figure 1bi)**. A comparable positional pattern of the vertex protein gene across multiple BMC types was observed **(Supplementary Table 1 and Figure 1b)**. The pentamer protein-encoding gene is found to be present relatively mid-operon positions. This recurring arrangement prompted us to question whether the gene’s location within the operon carries functional significance. In the PduBMC system, deletion of *pduN* yields leaky, tubular assemblies, confirming its role as a morphological keystone^14,17^. However, whether its operonic location inherently programs nanostructure assembly remained untested. Here, we treat gene order as an engineering variable to decode how genetic layout dictates nanoscale design. By repositioning *pduN* to multiple new locations within the *pdu* operon, we ask: Can operon rewiring direct the assembly of functionally distinct protein nanostructures? Our findings show that repositioning a single structural gene, without altering its coding sequence, can substantially alter nanocompartment morphology, enzyme encapsulation, and catalytic output. Together, these results indicate that operonic gene order represents an additional regulatory layer influencing the organization and morphological outcomes of BMC assembly. From a synthetic biology perspective, this highlights gene repositioning as a promising strategy to tune the architecture and function of protein-based nanodevices.

**Figure 1.**
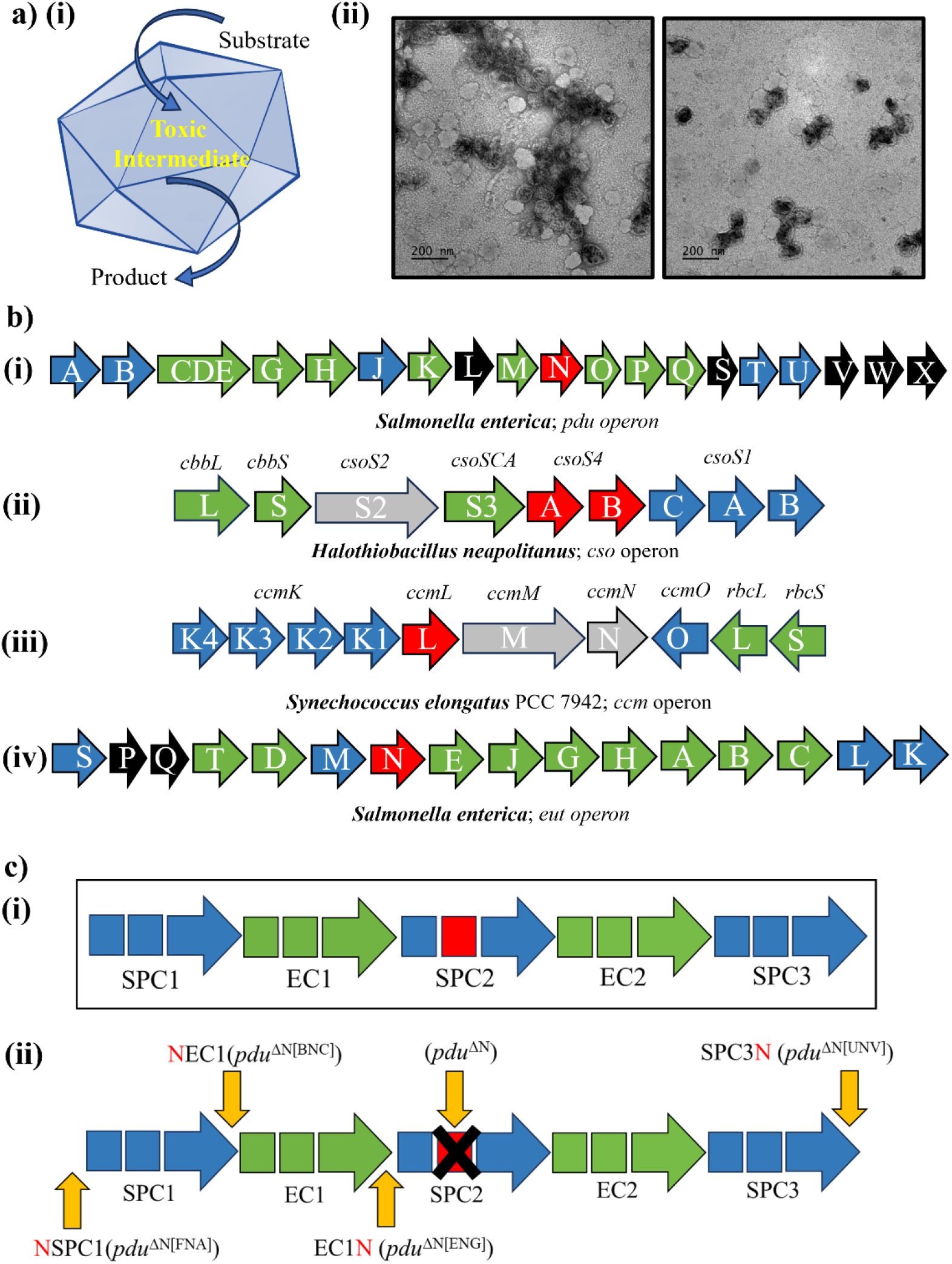
Structural organization of PduBMC and repositioning in *pdu* operon. General icosahedral morphology of PduBMC that sequesters toxic intermediate (propionaldehyde) by metabolizing 1,2 propanediol **ai)**, Transmission electron micrographs of purified PduBMC **aii)**, operons encoding different BMC systems including *pdu* operon, *cso* operon, *ccm* operon and *eut* operon **b)**, simplified *pdu* operon based on shell protein encoding gene cluster (SPC) and enzyme encoding gene cluster (EC) **ci)**, Highlighting repositioning of *pduN* generated in this study **cii)**.

## Materials and methods

### Chemicals and reagents

All chemicals are procured from Sigma-Aldrich unless mentioned. Agarose for gel electrophoresis is obtained from Lonza. B-PER II (Bacterial protein extraction reagent) was purchased from Thermo Fisher Scientific.

### Plasmids

pKD46, pMG1, and pLac22 plasmids used to perform recombination and transformation were a gift from Prof. Thomas A Bobik. The D-sfGFP plasmid was created using overlap extension PCR and cloned into the pLac22 plasmid.

### *In vivo* gene repositioning

Gene repositioning within the *pdu* operon was performed using the Datsenko and Wanner method based on λ Red recombination^18^. This scarless genetic modification was achieved by employing a dual-selection cassette, mPhes-Gent, wherein the mphes gene imparts sensitivity to 4-para-chloro-phenylalanine, and the gent gene confers resistance to gentamicin. The procedure was carried out in *Salmonella enterica* serovar Typhimurium LT2 harboring the pKD46 plasmid, which facilitates λ Red recombination. Initially, the *mPhes-Gent* cassette was inserted at the target site, followed by replacement with the PCR-amplified gene of interest. For precise integration, the forward primer (FP) is designed to include a homology arm corresponding to the sequence upstream of the insertion site, while the reverse primer (RP) includes a homology arm downstream of the gene of interest. During gene repositioning, care was taken to preserve the native ribosome binding site (RBS) sequences of both upstream and downstream genes. Successful recombination events are confirmed by PCR and Sanger sequencing (SS1001, SS1002, SS1003, SS1004, SS1005, SS1006, and SS1007).

### Expression and Purification of Pdu BMCs

To purify Pdu BMCs, a multi-step centrifugation and lysis protocol was followed^19^. Briefly, 100 µL of Salmonella enterica serovar Typhimurium LT2 wild-type and mutant strains were inoculated into 10 mL of LB medium supplemented with 0.6% (v/v) 1,2-propanediol (1,2-PD) and grown at 37 °C for 6 hours. Subsequently, 4 mL of this culture was used to inoculate 400 mL of NCE minimal medium supplemented with 0.6% 1,2-PD, 1 mM MgSO_4_, and 0.5% (w/v) succinate. Cultures were incubated for 14-16h at 37 °C with 180 rpm shaking. Cells were harvested by centrifugation at 7000 rpm for 10 minutes at 4ᵒC and washed twice with Buffer A (50 mM Tris-HCl, 500 mM KCl, 12.5 mM MgCl_2_, 1.5% 1,2-propanediol). The resulting cell pellet was resuspended in lysis buffer, consisting of Buffer A and B-PER II supplemented with 2.5 mg lysozyme, 3–5 mg DNase I, and 1 mM PMSF, and incubated at room temperature for 30 minutes on a rocker to facilitate lysis. Following lysis, cell debris was removed by centrifugation at 12,000 × g for 7 minutes twice at 4 °C. The clarified supernatant was further centrifuged at 20,000 × g for 20 minutes at 4 °C to pellet the BMCs. The resulting pellet containing BMCs was resuspended in Buffer B (50 mM Tris-HCl, 50 mM KCl, 5 mM MgCl_2_, 1% 1,2-propanediol) and stored or used for downstream applications.

### SDS-PAGE

For crude cell lysate, 3 mL of bacteria O.D. 1 was centrifuged, followed by washing with buffer A. The bacteria were lysed using buffer A, B-PERII, lysozyme, and PMSF with continuous shaking at room temperature for 30 mins. The cells were harvested by centrifuging at 12000 x g. The resulting 10µl clarified supernatant was loaded on an SDS-PAGE gel and stained with Coomassie. For Purified BMCs, 2µg protein sample was loaded and proceeded with silver staining.

### Transmission Electron Microscopy (TEM) Analysis

For TEM imaging, 10 µL of purified BMC sample at a concentration of 0.3 mg/mL was drop-casted onto a carbon-coated formvar mesh grid (Agar Scientific) and incubated for 3 minutes at room temperature. Excess sample was carefully removed using Whatman filter paper. The grid was then negatively stained with 10 µL of 0.6 mg/mL uranyl acetate for 45 seconds. Excess stain was blotted off, and the grid was briefly washed with syringe-filtered distilled water, followed by air-drying at room temperature. Prepared grids were imaged using a JEOL 2100 transmission electron microscope operated at 120 kV. Size measurements of the purified structures were performed using ImageJ software.

### Enzyme Activity Assay

The enzymatic activity of the PduCDE complex was measured using the MBTH (3-methyl-2-benzothiazolinone hydrazone hydrochloride) colorimetric assay^19^. For the assay, 5 µg of purified BMC was added to 950 µL of assay buffer composed of 0.2 M 1,2-propanediol, 0.05 M KCl, and 0.035 M potassium phosphate buffer (pH 8.0), and incubated at 37 °C. The reaction was initiated by the addition of 50 µL cyanocobalamin (15 µM final concentration). After a 10-minute incubation, the reaction was quenched by the addition of potassium citrate buffer (pH 3.6). Following quenching, 1 mL of 0.1% MBTH solution was added to the reaction mixture. After a 15-minute color development period, the absorbance of the reaction product was measured at 305 nm. The specific activity was calculated using a molar extinction coefficient of 13,000 M^−1^ cm^−1^. To assess PduCDE activity in crude lysates, the assay was performed using the same conditions, with 50 µL of clarified supernatant from lysed cells (explained in the SDS-PAGE section) substituted in place of purified BMC.

### RT-qPCR

A single colony of bacteria having different variants of *pdu* operons were inoculated into 10 mL of LB media.100 µL of overnight culture was inoculated in10 mL of LB medium supplemented with 0.6% (v/v) 1,2-propanediol (1,2-PD) and grown at 37 °C for 6 hours. Subsequently, 100µL of this culture was used to inoculate 10 mL of NCE minimal medium supplemented with 0.6% 1,2-PD, 1 mM MgSO_4_, and 0.5% (w/v) succinate for BMC expression. Total RNA was extracted using 1ml of 2hr,6hr and 14hr grown culture using SV Total RNA Isolation system (Promega corporation). RT-qPCR was performed using Go-Taq 1-step RT-qPCR reaction mix (Promega corporation) for *pduA* and *PduN* gene where 16S RNA was kept as control. One step RT-qPCR was performed using the protocol by standard kit using QuantStudio3 Real-Time PCR System (Thermo Fisher). To calculate the gene expression, fold change was calculated using CT value of each gene from all the variants.

### Growth Studies

Individual colonies of the wild-type and mutant strains were cultured overnight in LB medium at 37 °C with shaking at 180 rpm. The following day, the cultures were washed and diluted to an initial optical density (OD_600_) of 0.15 in no-carbon essential (NCE) minimal medium, supplemented with 3 mM amino acids (valine, leucine, isoleucine, and threonine), 1 mM histidine, 0.6% 1,2-propanediol,1 mM MgSO_4_ and 20nM cyanocobalamin to induce BMC formation. Culture samples were aliquoted every 6 hours, followed by OD_600_ determination and aldehyde estimation.

### Aldehyde detection

100µl potassium citrate buffer and 50µl of 0.1% MBTH were added to 100µl of cell lysate observed from all the bacterial strains. The reaction mixture was incubated for 10 minutes at 37ᵒC. The reaction was quenched using 100µl distilled water, followed by detection of the product at 305nm.

### Field Emission Scanning Electron Microscopy (FESEM) Analysis

For morphological analysis, 10 µL of overnight cultures of OD_600_ 0.1) of each mutant strain were drop-cast onto ultra-clean silicon wafers and allowed to air dry at room temperature. The dried samples were then imaged using field emission scanning electron microscopy (FESEM) at an accelerating voltage of 30 kV.

### Confocal microscopy

After confirming successful electroporation of the D-sfGFP construct into all mutant strains, the bacteria were cultured in BMC-inducing medium supplemented with 10 µM IPTG. A 10 µL aliquot of each culture of O.D_600_ 1 was fixed onto a glass slide and subsequently examined using a Zeiss LSM880 Airyscan microscope under both brightfield and FITC fluorescence channels.

## Results

### Rationale and Strategy for Gene Repositioning

Genetic programming of self-assembling nanostructures requires precise control over component expression and interactions^7,20^. While operon organization is known to regulate gene expression^2,21,22^, its role as a determinant of nanoscale architecture, governing the size, shape, and functional output of protein-based nanocompartments, remains unexplored. Notably, *pduN* occupies a mid-operon position within the *pdu* operon (Figure 1b), raising the possibility that its genomic location may influence assembly processes. Based on this observation, we hypothesised that gene position encodes structural and functional instructions for nanoscale morphogenesis. To test this, we treated the operon sequence as an engineerable variable and systematically repositioned *pduN* within the *pdu* operon. The following four variants were generated **(Figure 1cii and Supplementary Figure 1b)**: (i) upstream of primary shell protein gene *pduA* [*pdu*^ΔN[FNA]^], (ii) preceding adjacent to the primary signature enzyme *pduCDE* [*pdu*^ΔN[BNC]^], (iii) downstream of *pduCDE* [*pdu*^ΔN [ENG]^], and (iv) at the operon terminus following all shell protein genes [*pdu*^ΔN[UNV]^]. All variants were constructed using λ-Red recombination^18^ methodology with selection mediated by the mPhes-gent cassette^23,24^. These constructs were introduced into a deleted *pduN* (*pdu*^ΔN^) strain to isolate the impact of native gene position on BMC morphology, assembly, and metabolic activity. Wild-type (*pdu*^WT^) and *pdu*^ΔN^ individual strains were included as controls **(Supplementary Figure 2)**. This framework enabled us to systematically examine the influence of operonic architecture on bacterial organelle formation in a model compartment system.

### Influence of gene order on BMC Expression and Assembly

To determine whether repositioning of *pduN* altered the biogenesis or structural integration of nanocomponents, we analysed BMC-induced cell lysates and purified nanostructures from all engineered strains. Cultures were grown under minimal media conditions (NCE supplemented with 1,2 PD) that induce BMC formation, and lysates were subjected to compositional profiling using SDS-PAGE **(Supplementary Figure 3)**. All variants were observed to have protein bands with molecular weight similar to core shell protein PduBB′ and the catalytic enzyme complex PduCDE^25^, indicating that gene repositioning does not impair basic translation. This shows that operon rewiring retains expression of essential BMC constituents. We next assessed BMC assembly by purifying compartments from each strain and analysing their protein composition **(Figure 2)**. PduBB′ was consistently detected across all samples, suggesting robust shell formation regardless of *pduN* operon position **(Figure 2)**. Strikingly, enzyme encapsulation varied with *pduN* position in the operon. Notably, the *pdu*^ΔN[BNC]^ and *pdu*^ΔN[FNA]^ mutants lacked specific PduCDE bands in the purified BMCs. While in the other two mutants, *pdu*^ΔN[ENG]^ and *pdu*^ΔN[UNV]^, incorporation of PduCDE was observed as bands were visible. These results demonstrate that although the catalytic module, PduCDE, is synthesized in all mutants, its encapsulation is disrupted when *pduN* is placed upstream of or preceding the *pduCDE* gene, leading to failure in its integration to the BMCs. Further, some of the shell protein bands, such as PduA in *pdu*^ΔN[FNA]^ was missing, suggesting its non-incorporation within the purified BMC. Thus, the position of *pduN* within the operon determines effective BMC closure and consequently alters enzymatic cargo encapsulation. Simultaneously, the observed outcomes can be the result of early capping of the assembled BMC nanostructure that restricts the encapsulation.

**Figure 2.**
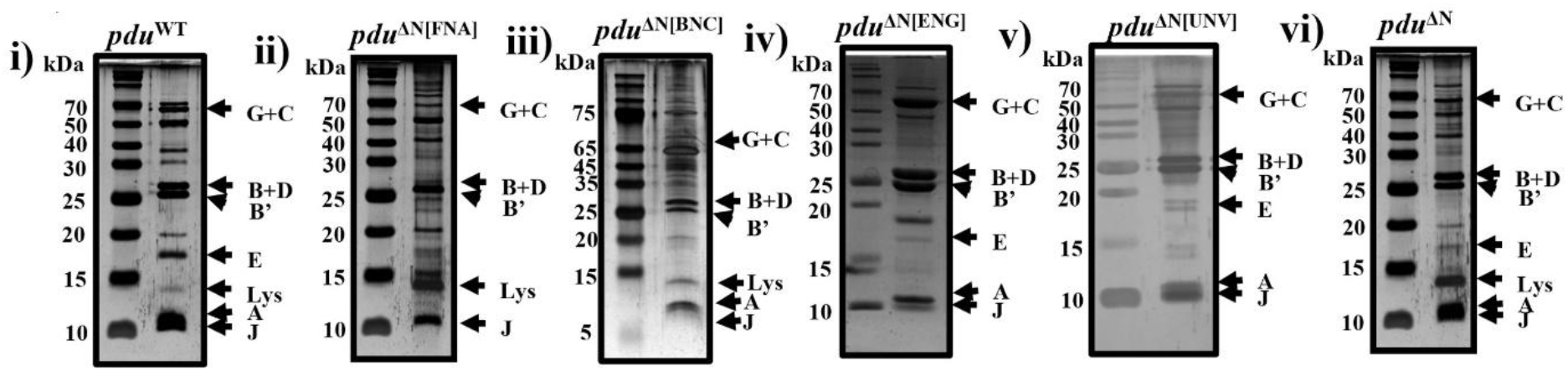
Expression of PduBMC native proteins. SDS-PAGE gel images of purified mutant BMCs, some of the major shell and enzyme protein bands are labeled to check the expression of BMC proteins

### Morphological Transitions in Repositioned BMC Assemblies

Transmission electron microscopy (TEM) of purified BMC variants revealed genetic repositioning of *pduN* gene reconfigures the nanoscale architecture **(Figure 3)**. Wild-type (*pdu*^WT^) PduBMCs formed uniform polyhedral structures ranging from 60–100 nm in diameter^13^ **(Figure 3ai,aii)**. In stark contrast, relocating *pduN* to the operon’s 5’-end, *pdu*^ΔN[FNA]^ induced aberrant self-assembly into discrete, spherical subunits (8–14 nm) **(Figure 3bi, bii)** often connected by fibrous extensions with diameters overlapping with the spherical structures **(Figure 3biii)**, suggesting that early operonic placement of *pduN* disrupts higher-order shell assembly and induces aberrant assembly of shell precursors. Also, the single shell proteins of PduBMC are well known for their self-assembling properties. Suggesting these fibrous extensions can be the result of the self-assembling morphology of single-shell proteins such as PduA/PduJ^26^. Further, positioning *pduN* preceding adjacent to *pduCDE* gene *pdu*^ΔN[BNC]^ yielded dense linear fiber bundles **(Figur3c)**. Interestingly, mutants with *pduN* placed downstream of *pduCDE* (*pdu*^ΔN[ENG]^ and *pdu*^ΔN[UNV]^) showed morphologically distinct tubular assemblies. *pdu*^ΔN[ENG]^ produced short tubes with irregular morphology (diameter: 40–80 nm; **Figure 3di,3dii**), while *pdu*^ΔN[UNV]^ formed long, tubes with wider diameter distribution (40–140 nm; **Figure 3ei, eii**). As reported in literature *pdu*^ΔN^ produced nanotubes^17^ (40-160nm; Figure 3fi,fii),. Collectively, these nanoscale phenotypes demonstrate that gene position orchestrates: (i) Assembly pathways, (ii) Structural fidelity, and (iii) Geometric precision.

**Figure 3.**
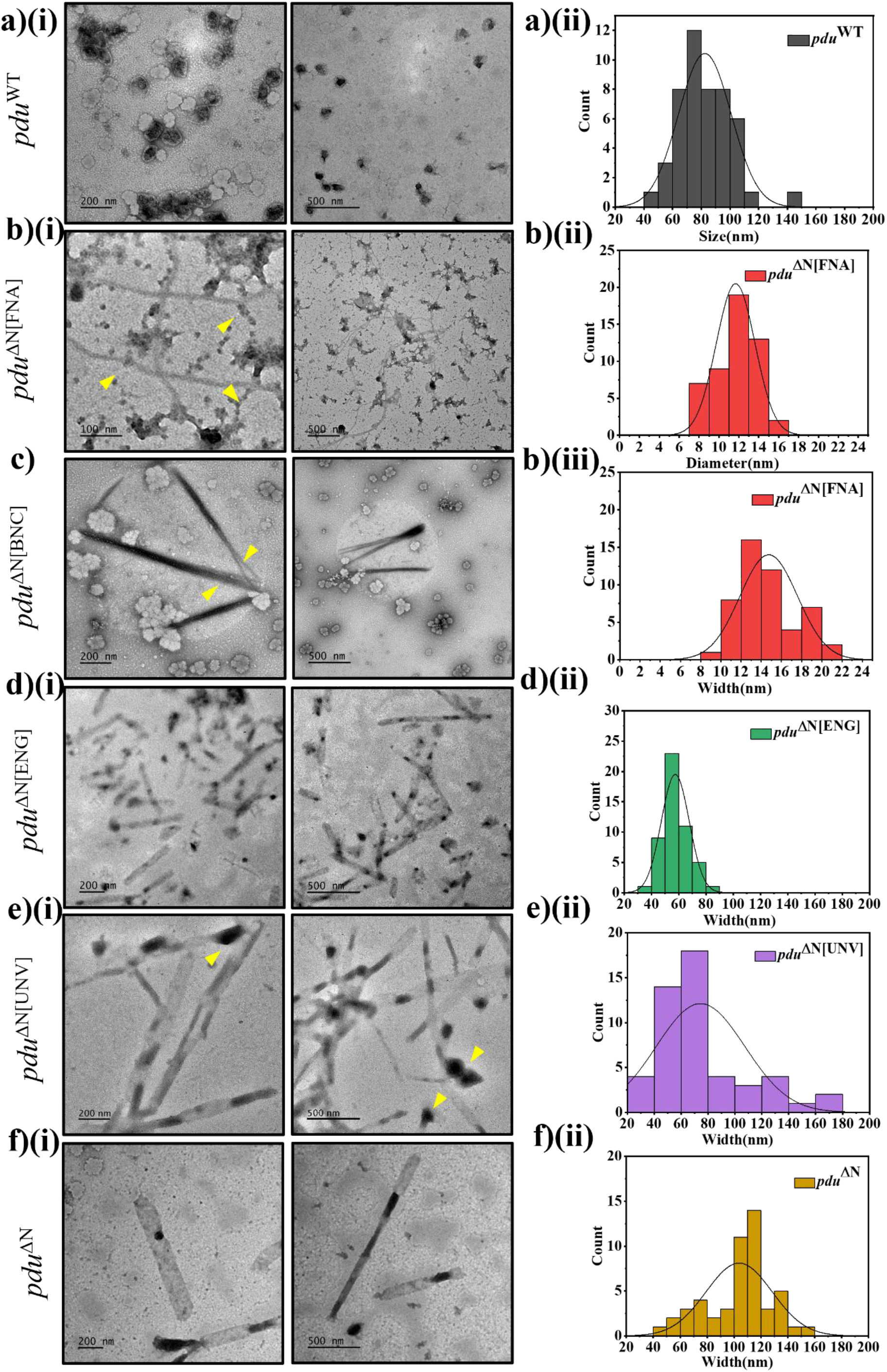
Morphological transitions in repositioned BMC assemblies. TEM images of BMCs purified from *pdu*^WT^ and the mutated operon expression. *pdu*^WT^ expressed as a polyhedral compartment **a)(i)** with a size ranging 60-100nm **a)(ii)**, *pdu*^ΔN[FNA]^ form small round structures **b)(i)** of diameter 8-14nm **b)(ii)** that assemble as fibers of overlapping width **b)(iii)**, *pdu*^ΔN[BNC]^ leads to formation of bundles of thin fibers **c)**, *pdu*^ΔN[ENG]^ assembled as short tube **d)(i)** of width 40-80nm **d)(ii)** and irregular morphological structures, *pdu*^ΔN[UNV]^ formed long tubes **e)(i)** with varying width from 40-140nm **e)(ii)**, As expected *pdu*^ΔN^ led to formation of tubes with of width 80-140nm **f)(i)(ii)**. The images are the representation of three independent experiments.

### BMC Cargo Loading and Functional Output

In SDS-PAGE data, we have observed a stark difference in the PduCDE bands with varying *pduN* position in the operon. To quantitatively analyze the functional consequences of nanostructures formed by the mutations, we quantified catalytic output and containment fidelity of PduCDE-loaded nanocompartments using an MBTH-based assay^27^ that monitors propionaldehyde production—a direct metric of repositioned BMC function **(Figure 4)**. In *pdu*^WT^, robust enzymatic activity was detected in both whole-cell lysates and purified BMCs, confirming the effective encapsulation and function of the enzyme.

**Figure 4.**
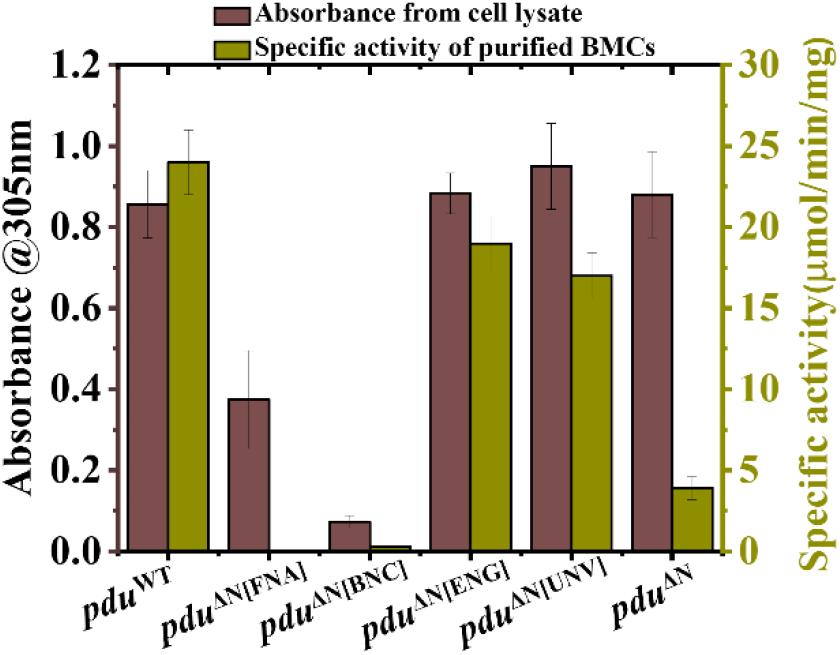
Enzyme encapsulation and functional consequences. The MBTH assay was conducted to assess PduCDE enzymatic activity using both whole-cell lysates and purified BMCs from various strains generated. While PduCDE activity was detectable in all lysate samples, it was absent in the purified BMC fractions of the *pdu*^ΔN[FNA]^ and *pdu*^ΔN[BNC]^ mutants, indicating defective enzyme loading. Data represented is the mean of three biological replicates (n = 3).

However, in mutants where *pduN* was repositioned upstream of *pduCDE, pdu*^ΔN[FNA],^ and *pdu*^ΔN[BNC]^, enzyme activity was present in lysates but undetectable in the purified BMCs. These findings align with SDS-PAGE data and indicate a severe impairment in enzyme encapsulation. This affected encapsulation can be due to the premature closure of the structure having *pduN* upstream of enzyme *pduCDE*. As PduN is reported to close the BMC structure and occupy the vertex at the last stage of BMC assembly^28^.

Notably, overall lysate activity was also diminished in these two strains, suggesting an impact on the protein expression levels, although quantification of level of expression is beyond this study and can be performed in future studies. In contrast, *pdu*^ΔN[ENG]^ and *pdu*^ΔN[UNV]^, where *pduN* was placed downstream of *pduCDE*, showed enzyme activity in both lysates and purified structures, confirming successful encapsulation. Enzymatic activity assay revealed that while *pdu*^ΔN[ENG]^ and *pdu*^ΔN[UNV]^ retained significant specific activity (19±1.7 and 17±1.4 µmol/min/mg, respectively), their activity was moderately reduced compared to *pdu*^WT^ (24±2.0 µmol/min/mg). As PduN is responsible for closing PduBMC. This decrease in activity likely reflects the reduction in packing of PduCDE within the BMC due to the *pduN* position. Such as the presence of *pduN* immediately downstream of *pduCDE* in *pdu*^ΔN[ENG]^ strain helps in closure with PduCDE more efficiently than *pdu*^ΔN[UNV],^ which has distantly placed *pduN*. Importantly, *pdu*^ΔN^ BMCs, despite showing wild-type levels of PduCDE activity in lysate, exhibited significantly lower activity in purified BMCs (3.9 ± 0.7 µmol/min/mg), reinforcing their characterization as “leaky tubes’’^29^. Together, these results demonstrate that *pduN* not only mediates shell closure, but its position also governs the efficient sequestration of enzymes. As we have seen, the position of *pduN* altered the BMC’s outer shell morphology (structure), which is a possible consequence of the alteration in shell protein packaging. So, the enzyme encapsulation can also be affected due to differences in the packing of cargo encapsulation mediating shell proteins, e.g. N-terminal of PduBB’^30^. This can be one of the major reasons behind this remarkably changed functional difference. Further, the BMC purification protocol involves differential centrifugation, preventing co-sedimentation of free enzyme proteins with BMCs, making this assay significant to check the encapsulation of enzyme proteins within the repositioned BMCs. Overall, our findings establish that the operonic positioning of *pduN* is a key determinant of both the morphological and functional integrity of Pdu BMCs. The native placement of *pduN* downstream of *pduCDE* likely represents an evolutionary optimization for coordinated shell closure and enzyme encapsulation.

### Reversing *pduN* position to *pdu*^WT^

Repositioning of *pduN* is found to alter both the structural and functional parameters of the PduBMC. To determine whether these effects arose specifically from *pduN* repositioning or are a consequence of genetic manipulation itself, we have performed reverse repositioning. In this approach, *pduN* was restored to its native genomic position in two of the generated strains (*pdu*^ΔN[BNC]^ and *pdu*^ΔN[ENG]^), followed by analysis of BMC structure and enzyme activity. The strains selected here are one upstream of the *pduCDE* gene and one downstream of it, where *pdu*^ΔN[BNC]^ mutant had both remarkable structural and functional transitions and *pdu*^ΔN[ENG]^ had major structural transitions. SDS–PAGE analysis of purified BMCs from the reverse-positioned *pdu*^ΔN[BNC]^ and *pdu*^ΔN[ENG]^ strains revealed the presence of all shell proteins and encapsulated enzymes, with banding patterns comparable to those observed for *pdu*^WT^ **(Figure 5a)**.

**Figure 5.**
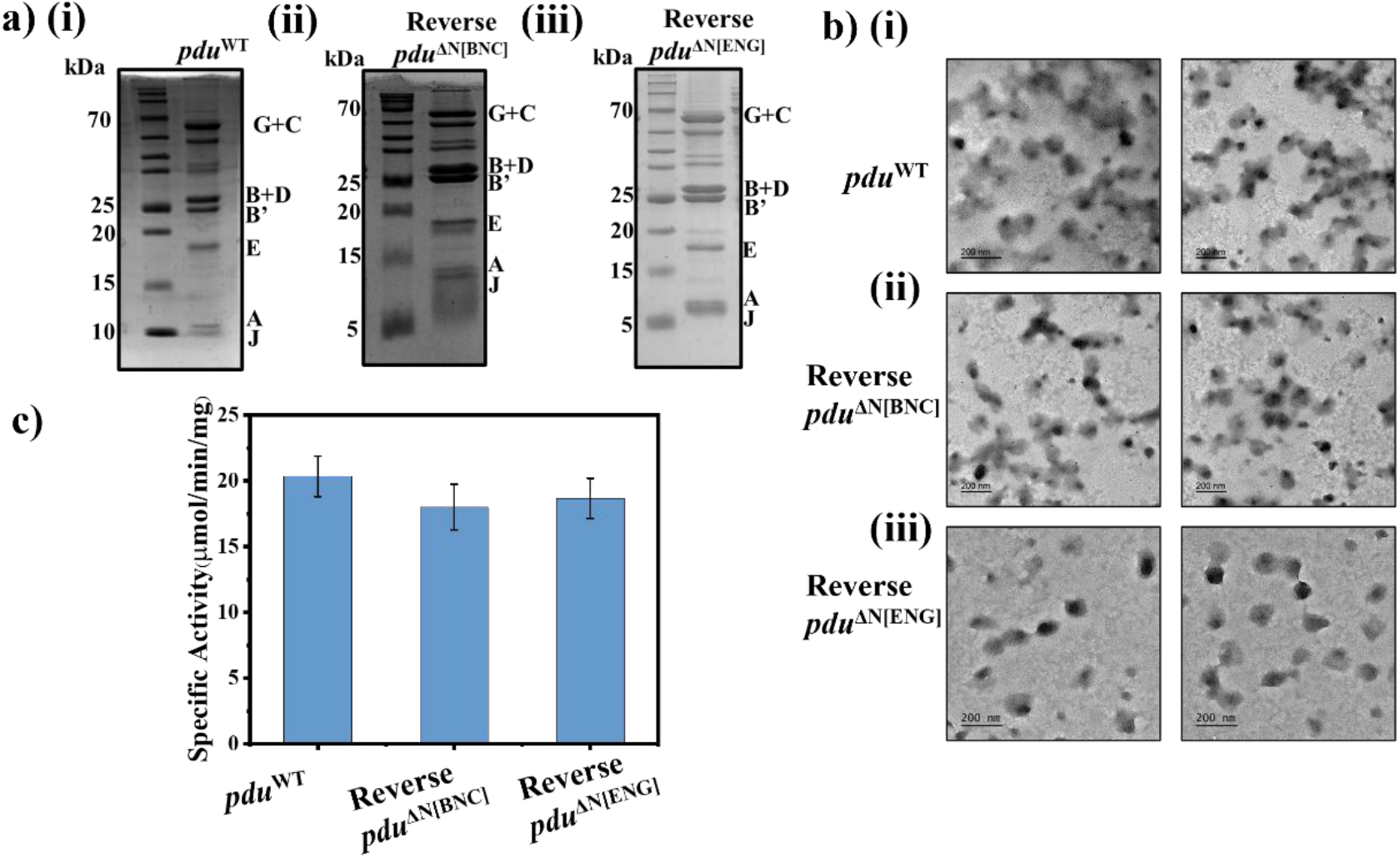
Effect of reversing *pduN* positioning on bacterial microcompartment structure and function. SDS-PAGE images of purified WT and reversed strain BMCs, having bands for all shell and enzyme proteins **a)**, TEM images of purified WT and reversed strain BMCs showing polyhedral structure **b)**, The MBTH assay was conducted to assess PduCDE enzymatic activity using purified BMCs from the reversed strain generated. PduCDE activity was detectable in all samples, indicating successful enzyme loading **c)**. Data represented is the mean of three biological replicates (n = 3).

A notable difference was observed in the original *pdu*^ΔN[BNC]^ strain, where prominent bands corresponding to enzymes were absent (Figure 2biii). Consistent with these findings, TEM analysis showed that the fibrous structures formed by *pdu*^ΔN[BNC]^ and the short tubular structures formed by *pdu*^ΔN[ENG]^ (Figure 3c and 3e) reverted to the native polyhedral PduBMC morphology upon restoration of *pduN* to its native position **(Figure 5b)**. The presence of enzyme protein bands in SDS–PAGE indicated retention of enzymes within the purified reverse strains. We have further validated this observation by the MBTH enzyme activity assay, which demonstrated enzyme activities comparable to *pdu*^WT^ in both reverse-positioned strains **(Figure 5c)**. Together, these results confirm that although *pdu*^ΔN[BNC]^ and *pdu*^ΔN[ENG]^ mutants had different structures and enzyme activity, both reverted to the *pdu*^WT^ phenotype structurally and functionally upon restoration of *pduN* native positioning. Overall, this study establishes that the observed alterations in BMC architecture and function are specifically driven by *pduN* positioning rather than by nonspecific effects of genetic manipulation. Further to understand the mechanistic connection between gene order and assembly dynamics of PduBMC, we have performed a time-dependent RT-qPCR study. While RT-qPCR results cannot directly infer stoichiometry between transcript levels and shell incorporation, the data indicate that operon rewiring alters mRNA expression dynamics as a function of gene position, correlating with distinct assembly outcomes **(Supplementary Figure 4)**. The differences in mRNA levels can be due to its differential degradation across the operon, there is transcriptional stalling, or, given the size of the operon, the changes to the operon are causing changes to mRNA secondary structure that are affecting the efficiency of cDNA synthesis.

### Bacteria Viability as a Metric for Nanoreactor Performance

The designed repositioned BMCs may be used as functional bioreactors that must operate compatibly within living hosts. As mentioned, PduBMC is involved in the metabolic functioning of bacteria by restricting the escape of toxic intermediate, propionaldehyde. The efficient assembly and operation of BMC is critical for sustaining bacterial viability in their native nutrient-limited environment. So, the effect of gene repositioned BMC structure and enzymatic encapsulation on the cellular fitness of all the mutants was evaluated. All the strains were cultured in minimal media (NCE) containing 1,2 PD as the sole carbon source, supplemented with limiting (20nM) **(Figure 6a,6b)** concentrations of cyanocobalamin (cyano-B_12_) to support cofactor-dependent PduCDE activity. Under limiting conditions, the *pdu*^ΔN[FNA]^ mutant displayed no detectable initial growth, but grew rapidly with time. The *pdu*^ΔN[BNC]^ strain exhibited faster growth, with a significant specific growth rate of 0.015h^−1^ compared to *pdu*^WT^ 0.004h^−1^. This growth kinetics likely stems from free cytosolic PduCDE activity that permits metabolism of 1,2 PD. The growth with only 1,2 PD as a carbon source proves the presence of limited cellular PduCDE availability. Other mutants where *pduN* is repositioned downstream of *pduCDE pdu*^ΔN[ENG]^, *pdu*^ΔN[UNV]^, and *pdu*^ΔN^ mutants showed a growth profile faster than the wild-type, with a specific growth rate of 0.007h^−1^, 0.023h^−1^, and 0.028h^−1^, respectively. This growth acceleration is attributed to increased cytosolic enzyme availability resulting from partial encapsulation, corroborated by specific activity measurements. Together, these data strongly support that mispositioning *pduN* upstream of *pduCDE* disrupts proper enzyme encapsulation, leading to functional impairment. Conversely, downstream positioning compromises encapsulation efficiency due to altered BMC architecture but retains partial metabolic functionality. These findings highlight a direct link between gene order, metabolic compartmentalization, and organism fitness. Such phenotypes underscore the evolutionary pressure to maintain operon configurations that ensure efficient encapsulation and detoxification in environments rich in 1,2 PD, a common nutrient in the mammalian gut^31,32^.

**Figure 6.**
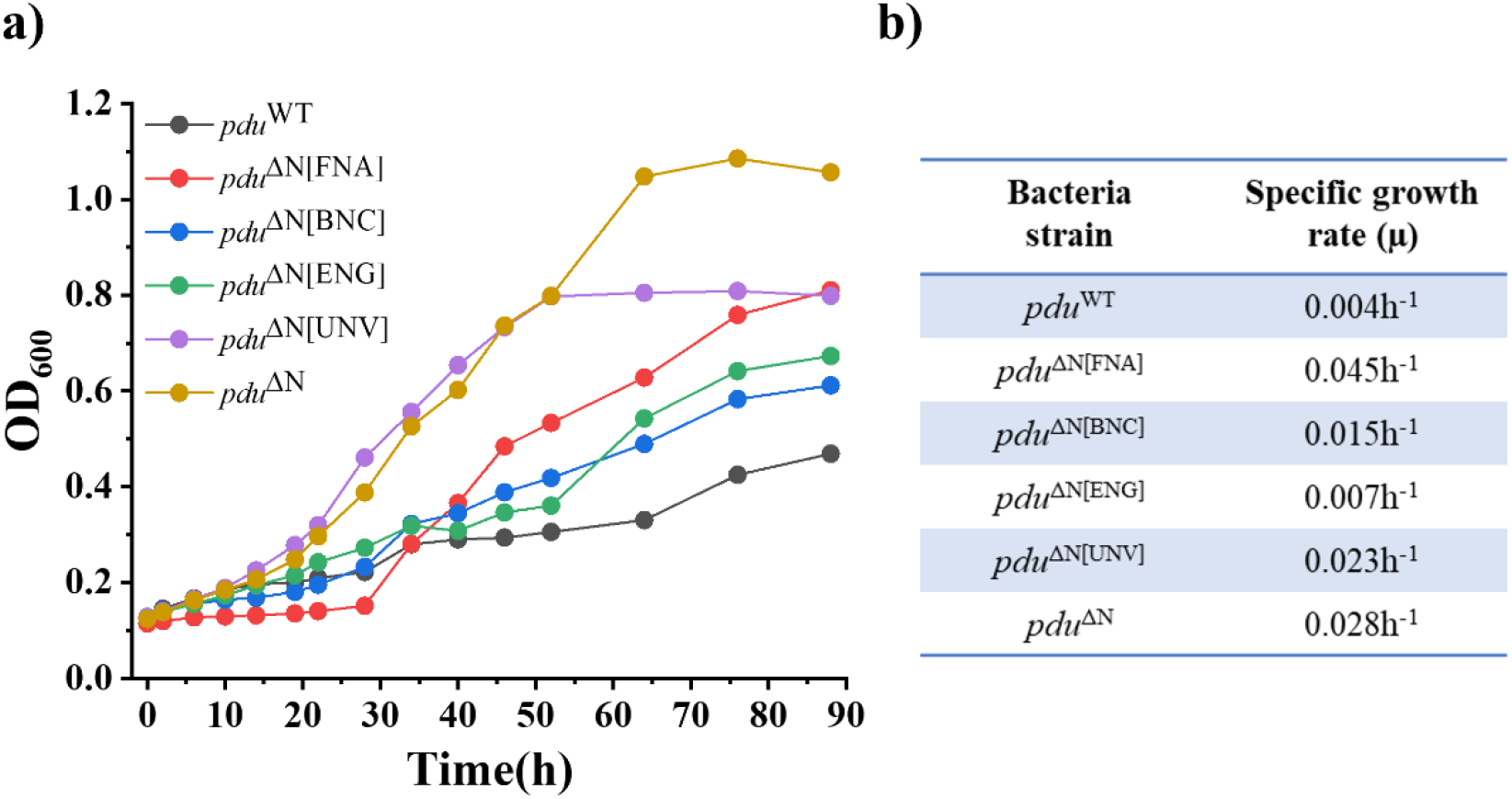
Effect of bacterial microcompartment function on cellular fitness. Bacterial growth was assessed in minimal media under minimal media having 1,2-propanediol (1,2 PD) as the sole carbon source, with limiting (20nM) concentrations of cyanocobalamin (cyano-B_12_). All the mutants were growing based on the free and encapsulated enzyme (PduCDE) activity, such that the more the free enzyme activity, the faster the growth rate **a).** The specific growth rate for each mutant strain was determined based on log phase growth kinetics **b)**. The growth curves are representative of three independent experiments.

### Aberrant BMC Structures Alter Bacterial Morphology

Successful integration of engineered nanostructures requires physical compatibility with the host chassis. Field emission scanning electron microscopy (FESEM) revealed that misassembled repositioned BMCs induce profound ultrastructural defects in bacterial cells **(Figure 7)**. *pdu*^WT^ exhibited native rod-shaped morphology **(Figure 7ai)**. Whereas, cells harboring the *pdu*^ΔN[FNA]^ construct displayed pronounced morphological distortion, with irregular, deformed envelopes and collapsed structures **(Figure 7aii)**, directly linking to cytotoxic effects of intracellular aldehyde accumulation, likely inducing oxidative stress and damaging macromolecular structures, including membrane components^33^. In the *pdu*^ΔN[BNC]^ mutant, FESEM imaging revealed intercellular connectivity, forming undivided chains of cells **(Figure 7aiii)**. These morphological defects align with prior TEM observations of intracellular fibrous aggregates and suggest that misassembled shell components interfere with cytokinesis, possibly through steric obstruction^34,35^. Cells expressing the *pdu*^ΔN[ENG]^ operon, due to the formation of short tubular BMCs, exhibited similar morphology to *pdu*^WT^ **(Figure 7aiv)**. Strikingly, both *pdu*^ΔN[UNV]^ and *pdu*^ΔN^ strains form long tubular BMCs that display highly ordered, chain-like arrangements of cells **(Figure 7av and 7avi)**. These linear alignments suggest partial or complete inhibition of cell fission, likely due to elongated internal structures traversing the cytoplasm and obstructing septation. To check this, we compared the length of WT bacteria with the purified BMC structure lengths **(Sup Figure 4)**. The comparative data showed that the size of *pdu*^WT^ BMC and *pdu*^ΔN[ENG]^ BMC structures was less than the entire cell, whereas the lengths of *pdu*^ΔN[BNC]^, *pdu*^ΔN[UNV]^, and *pdu*^ΔN^ BMCs surpassed the bacteria’s size. Thus, enabling these structures to obstruct the cytokinesis. In literature, cellular obstruction has been observed in the PduBMC system when tubular structures are formed, such as overexpression of some shell proteins, such as PduA, PduJ, PduABB’^26,35,36^ in a heterologous host, and deletion of genes *pduN* and pduJ^14,28^ within the native operon. In contrast to those studies our work reports that switching the position of the gene within the operon of the native host influences BMC formation at the cellular level. The consistency of this phenotype across *pdu*^ΔN[BNC]^, *pdu*^ΔN[UNV]^, and *pdu*^ΔN^ distinct constructs underscores the mechanical impact of extended cellular architecture. It is worth mentioning here that the ectopic expression of *pduN* rescued these tubular formations in the *pdu*^ΔN^ mutant, but its expression from different positions behaved dramatically^17^. Further, interference in cytokinesis and cellular spatial organization due to these elongated BMC tubes supports the idea that natural selection has favoured a gene order that ensures not just expression but also the physical compatibility of shell assembly within the crowded intracellular environment^36^. Among all mutants, only *pdu*^ΔN[FNA]^ displayed the distorted morphology suspected due to the free enzyme and aldehyde. To confirm if the change was due to aldehyde, we checked the cellular aldehyde levels using the MBTH assay. The resulting data confirm the presence of the highest level of aldehyde in this mutant, which supports our observation **(Figure 7c)**. This data again highlights the importance of PduBMC in the accumulation and conversion of toxic aldehyde, providing the bacteria a growth advantage^33,37^. Thus, disruption of canonical BMC assembly leads to altered cell shape, impaired division, and stress-induced deformation. These findings emphasize the broader cellular consequences of organelle misassembly and highlight the critical role of native gene positioning in maintaining both metabolic and morphological homeostasis.

**Figure 7.**
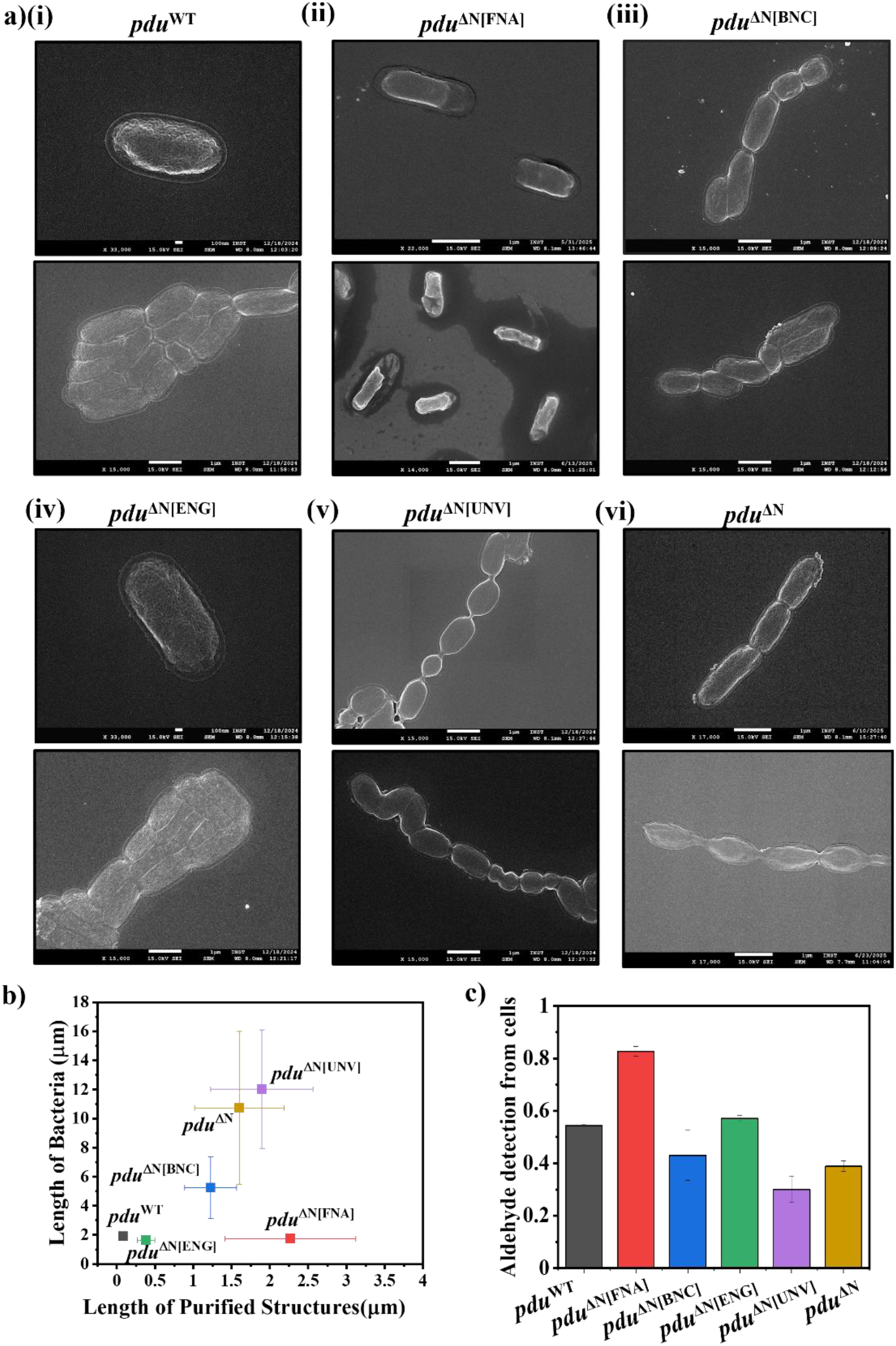
Effect of bacterial microcompartment structure and function on cellular morphology. FESEM imaging was used to investigate morphological alterations across various strains. The *pdu*^WT^ displayed normal rod-shaped morphology and cell clustering **ai)**, while the *pdu*^ΔN[FNA]^ mutant exhibited a distorted cellular shape **aii)**, *pdu*^ΔN[ENG]^ strain had similar morphology to *pdu*^WT^ **aiv)**, In contrast, longitudinally connected chains of cells were evident in the *pdu*^ΔN[BNC]^ **aiii)**, *pdu*^ΔN[UNV]^ **av)**, and *pdu*^ΔN^ **avi)** mutants, suggesting aberrant division phenotypes. The images displayed here are the representative images for 3-5 independent experiments. A comparison of the purified BMC length and bacterial length illustrates their direct relation **b)**. The data compared here is the mean of lengths measured using ImageJ. To assess aldehyde accumulation, bacterial cultures grown with both succinate and 1,2 PD were harvested and subjected to the MBTH-based aldehyde detection assay. The highest level of aldehyde is detected in *pdu*^ΔN[FNA]^ mutant **c)**. The data represented is the mean of three biological replicates (n = 3).

### Tracking of BMC Structure Within Bacteria

Following the assessment of how *pduN* repositioning influences BMC structure, function, and bacterial morphology, we investigated the subcellular localization of the BMCs using a fluorescence-based approach. To this end, the N-terminal 20 amino acids of the PduD enzyme, known as the encapsulation peptide, were fused to superfold GFP (sfGFP), generating the reporter construct D-sfGFP. This peptide is recognized for its specific interaction with the shell protein PduBB′, thereby facilitating targeted interaction with the PduBMC^30,38,39^. Fluorescence microscopy of the *pdu*^WT^ expressing D-sfGFP revealed cytoplasmic punctate fluorescence, indicative of a uniform distribution of BMCs throughout the cell **(Figure 8ai)**. In contrast, the mutant strain *pdu*^ΔN[FNA]^ displayed distinct fluorescence puncta localized predominantly at the poles of the cells, suggesting the presence of aberrantly BMCs at the cell ends **(Figure8aii)**. This polar localization corroborates our TEM observations and supports the formation of malformed BMC architecture upon this *pduN* repositioning. The *pdu*^ΔN[BNC]^ mutant, characterized by connected cellular morphology, exhibited a more diffuse fluorescence pattern throughout the cytoplasm **(Figure 8aiii)**. As this mutant was observed to have free enzyme PduCDE. This diffused pattern may result from the restricted encapsulation of D-sfGFP within BMCs. Further, in the mutants where *pduN* was repositioned downstream within the operon (*pdu*^ΔN[ENG]^ and *pdu*^ΔN[UNV]^), distinct patterns of fluorescence were observed, aligning with morphological features noted via FESEM. Specifically, *pdu*^ΔN[ENG]^ exhibited a centralized accumulation of fluorescent signal within cells **(Figure 8aiv)**, while *pdu*^ΔN[UNV]^ displayed longitudinal fluorescence extending along chains of connected bacterial cells **(Figure 8av)**. This longitudinal fluorescence provides visual evidence of enzyme encapsulation in *pdu*^ΔN[UNV]^ mutant, as corroborated by SDS-PAGE and MBTH assay data. A similar longitudinal fluorescence distribution was also evident in the *pdu*^ΔN^ control mutant **(Figure 8avi)**. Collectively, these observations confirm the successful tracking of BMCs using D-sfGFP and demonstrate that *pduN* positioning significantly affects both BMC localization and assembly dynamics within the bacterial cytoplasm, aligning with the FESEM data.

**Figure 8.**
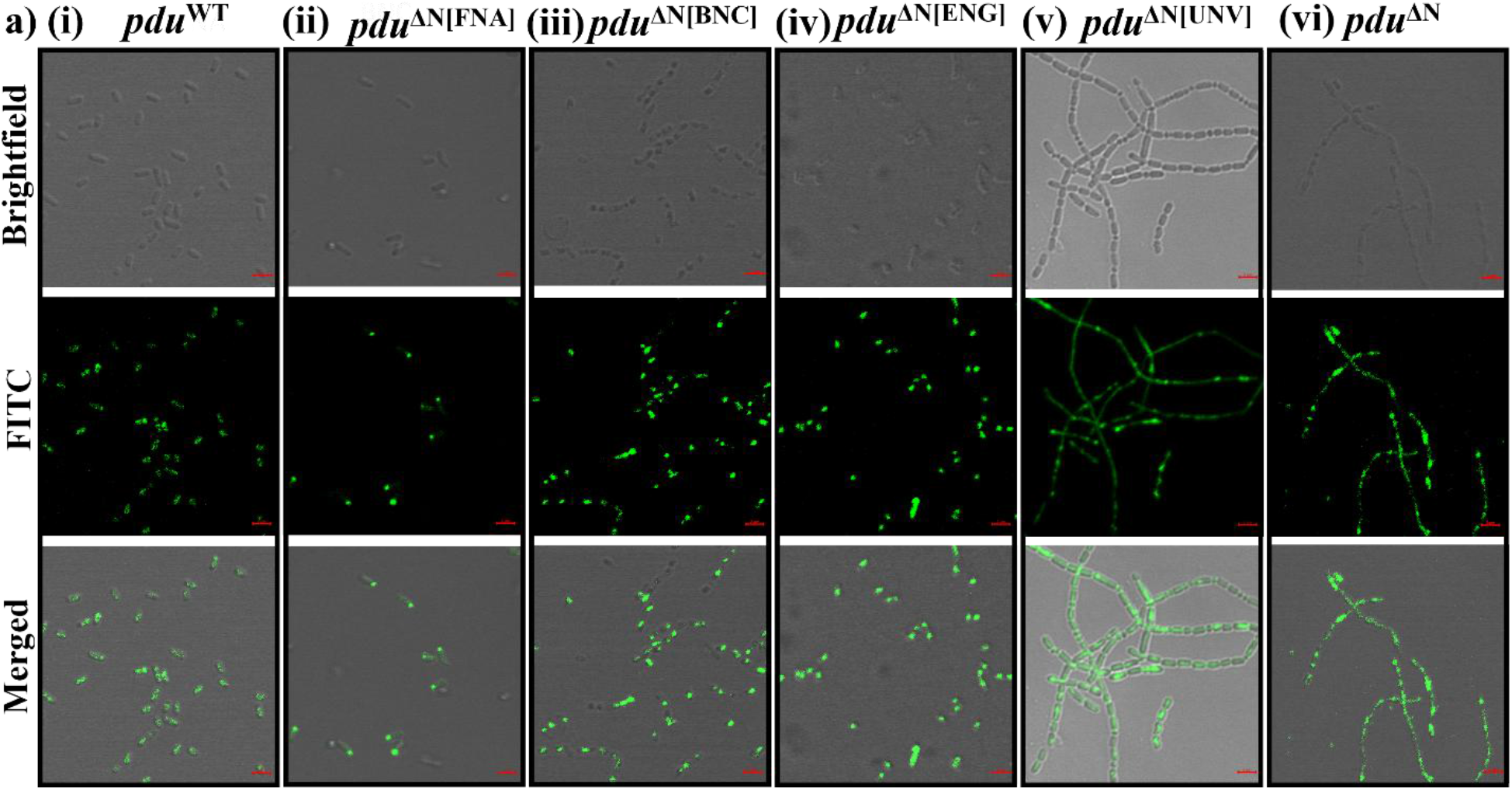
Tracking of BMC Structure Within Bacteria. Confocal imaging was performed to examine the subcellular distribution of BMCs in WT and mutant strains expressing D-sfGFP. Distinct fluorescent puncta were distributed throughout the cytoplasm in the *pdu*^WT^ strain **ai)**, in the *pdu*^ΔN[FNA]^ mutant, fluorescence localized predominantly at the bacterial poles **aii)**, the *pdu*^ΔN[BNC]^ mutant exhibited intercellular connections characterized by continuous fluorescent puncta **aiii)**, centralized fluorescence clusters were observed in the *pdu*^ΔN[ENG]^ mutant, indicating a compact BMC distribution **aiv)**, in both *pdu*^ΔN[UNV]^ and *pdu*^ΔN^ mutants, longitudinally connected cells with traversing fluorescent signals were detected **av, avi)**. The images displayed here are representative of 3-5 independent experiments. All images were captured at a 2 µm scale.

## Discussion

Bacterial microcompartments offer a unique platform to study genetically controlled protein-based organelles that can compartmentalize specific metabolic activities and perform different reaction kinetics. Our findings suggest that successful engineering of such systems must consider not only the gene expression but also their relative operonic positions from where they are expressed. Existing literature has explored the expression through gene deletion strategy, resulting in differences in protein expression, BMC morphology, and enzyme encapsulation, followed by differences in BMC activity^14,19,35^. Another study shows how the position of *pduJ* altered its function when placed at the position of *pduA*^40^. These studies explored the function of each of the shell protein in the assembly and function of PduBMC. Here, in this study we extended our question on gene position of the only pentameric protein-encoding gene by placing it at different positions throughout the *pdu* operon. Our study showed that though there is no complete deletion of the gene (*pduN*), the resultant BMC expression, morphology, encapsulation and activity were altered. Initial electron microscopy analyses reveal that *pduN* relocation reprograms self-assembly in such a way that early operonic placement induces BMC into disordered spherical structures/fibers, while terminal positioning enables delayed capping into extended nanotubes—contrasting sharply with evolutionarily optimized polyhedra at native loci **(Fig. 3)**. This establishes gene order as a regulator of hierarchical assembly. Crucially, spatial adjacency between *pduN* and enzymatic genes *pduCDE* governs cargo-loading fidelity: upstream positioning causes complete containment failure, whereas distal placement allows enzyme packing as observed in SDS-PAGE images and MBTH assay data **(Fig. 2 and Fig.4)**. These effects were further validated using a reverse repositioning study **(Fig.5)**, where placing *pduN* back to its native place was observed to rescue PduBMC morphology and enzyme activity. Thus, confirming the changed structural and functional properties of PduBMC due to *pduN* repositioning. Thus, our study demonstrated that *pduN’s* operonic location encodes both geometric control and functional coordination for the protein-based nanoreactors. As the functional aspect of BMCs is directly correlated to their metabolic requirement, we extended to analyse the growth rate of each of the mutants, which showed that the nanocompartment structure can control the growth rate of bacteria **(Fig.6)**, demonstrating that gene repositioning balances metabolic flux with containment—a prerequisite for biosafe nanoreactor design. This mechanistic understanding enables rational design strategies to optimize flux control, minimize cytotoxic intermediates, and ensure host viability^37,41^. Along with the structural and functional control, our data reveal that aberrant nanostructures exceeding cellular dimensions (>1.5 μm) physically obstruct cell division, while optimized geometries (polyhedral/short tubes) maintain ultrastructural integrity. Further, the dimensions of nanostructures have been observed to control the lengths of connected bacteria **(Fig. 7b)**. This makes these nanostructures a promising tool to utilize genetically controlled systems for customized applications. Other than serving as a tool for engineering BMCs for applications, this study provides support to the fundamental study of operon encoded assembly logic. Literature suggests that the operon order reflects the order of protein complex assembly ^22^and gene arrangement holds information about an evolutionarily optimized problem^42^. These comparative and modelling studies predicted gene-order–driven assembly patterns, where the direct experimental validation has been limited. Our results provide empirical evidence supporting this hypothesis, showing that altering gene order alone is sufficient to influence the assembly and final architecture of a BMC. Aligning with the gene order-driven logic, our study provides the evolutionary significance of the middle position of *pduN* within the *pdu* operon. We have found that the *pduN* position controls PduBMC in such a way that upstream *pduN* repositioning leads to impairment in enzyme encapsulation, thus leading to aldehyde toxicity within bacteria.

Whereas distant downstream positioning prevents cytokinesis, another essential aspect for bacterial survival **(Figure 9)**. Thus, *pduN* positioning carries evolutionary information regarding PduBMC assembly as well as its importance at the cellular level to carry out its function efficiently.

**Figure 9.**
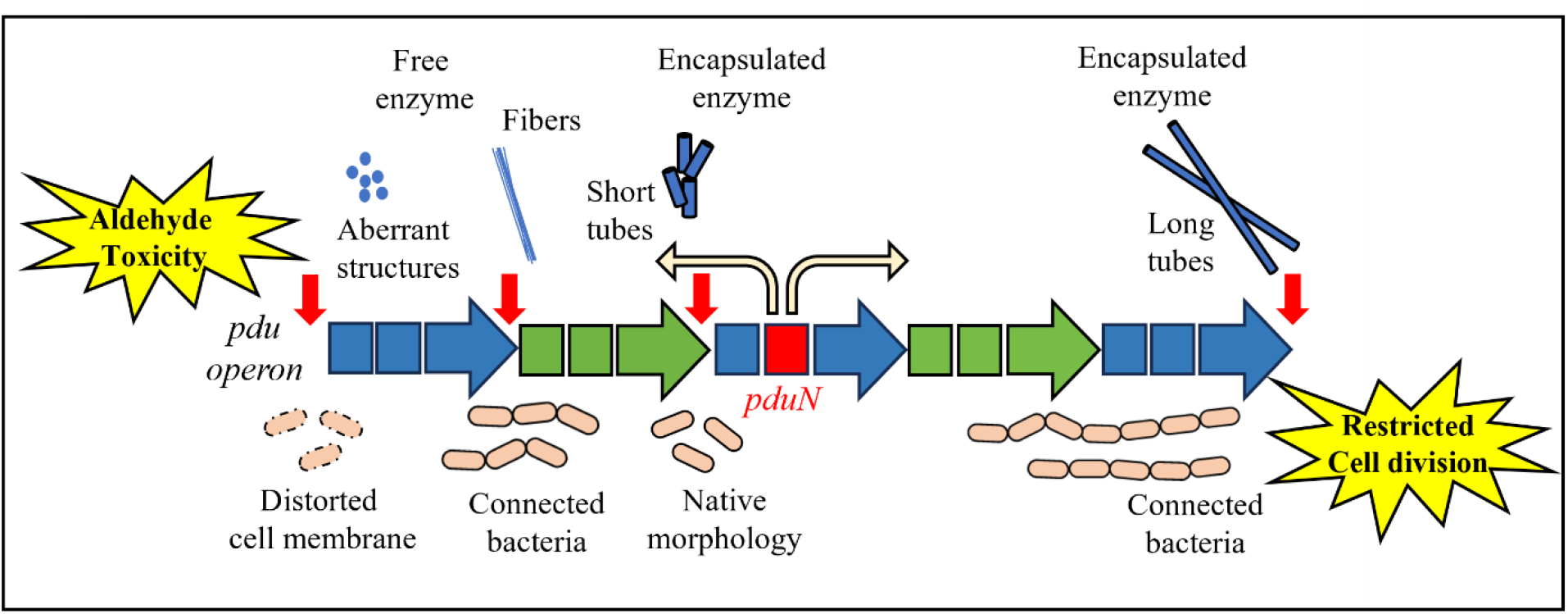
Schematic representation of structural and physiological outcomes of *pduN* repositioning.

## Conclusion

Our study positions gene order as a fundamental genetic framework that orchestrates assembly and drives nanoscale morphogenesis. By repositioning a single structural gene, *pduN*, we reprogrammed the size, geometry, and functional output of self-assembling bacterial microcompartments, demonstrating that operon architecture encodes both structural and functional logic for hierarchical protein assembly. By decoding how genetic syntax directs nanoscale self-assembly, we advance an experimental framework for building protein-based nanodevices *in vivo* and *in vitro*. Our findings also underscore the evolutionary significance of gene order in regulating BMC assembly and promoting bacterial fitness and survival.

## Author’s contribution

DG designed and conducted the experiments, analysed the data, and wrote the manuscript draft. PN and AR conducted experiments and analysed data. SS conceived the idea, designed the experiments, and wrote the manuscript.

## Acknowledgments

The authors acknowledge INST for the central instrument facilities. DG acknowledges CSIR India for the fellowship, PN acknowledges UGC India for the fellowship, and AR acknowledges INST for the fellowship. The authors acknowledge SS lab members for valuable discussion.

## References

(1) Lim, H. N.; Lee, Y.; Hussein, R. Fundamental Relationship between Operon Organization and Gene Expression. Proceedings of the National Academy of Sciences 2011, 108 (26), 10626– 10631. 10.1073/pnas.1105692108.

(2) Chizzolini, F.; Forlin, M.; Cecchi, D.; Mansy, S. S. Gene Position More Strongly Influences Cell-Free Protein Expression from Operons than T7 Transcriptional Promoter Strength. ACS Synth. Biol. 2014, 3 (6), 363–371. 10.1021/sb4000977.

(3) Zhang, W.; Ren, D.; Li, Z.; Yue, L.; Whitman, W. B.; Dong, X.; Li, J. Internal Transcription Termination Widely Regulates Differential Expression of Operon-Organized Genes Including Ribosomal Protein and RNA Polymerase Genes in an Archaeon. Nucleic Acids Research 2023, 51 (15), 7851–7867. 10.1093/nar/gkad575.

(4) Sáenz-Lahoya, S.; Bitarte, N.; García, B.; Burgui, S.; Vergara-Irigaray, M.; Valle, J.; Solano, C.; Toledo-Arana, A.; Lasa, I. Noncontiguous Operon Is a Genetic Organization for Coordinating Bacterial Gene Expression. Proceedings of the National Academy of Sciences 2019, 116 (5), 1733–1738. 10.1073/pnas.1812746116.

(5) Pai, D. A.; Engelke, D. R. Spatial Organization of Genes as a Component of Regulated Expression. Chromosoma 2010, 119 (1), 13–25. 10.1007/s00412-009-0236-2.

(6) Ma, Q.; Xu, Y. Global Genomic Arrangement of Bacterial Genes Is Closely Tied with the Total Transcriptional Efficiency. Genomics Proteomics Bioinformatics 2013, 11 (1), 66–71. 10.1016/j.gpb.2013.01.004.

(7) Bedi, S.; Rose, S. M.; Kaur, S.; Negi, P.; Sinha, S. Protein-Protein Interactions as Determinants of Operon Architecture. Biochimica et Biophysica Acta (BBA) - General Subjects 2025, 1869 (6), 130794. 10.1016/j.bbagen.2025.130794.

(8) Macnab, R. M. How Bacteria Assemble Flagella. Annu Rev Microbiol 2003, 57, 77–100. 10.1146/annurev.micro.57.030502.090832.

(9) Diepold, A.; Armitage, J. P. Type III Secretion Systems: The Bacterial Flagellum and the Injectisome. Philos Trans R Soc Lond B Biol Sci 2015, 370 (1679), 20150020. 10.1098/rstb.2015.0020.

(10) Yeates, T. O.; Crowley, C. S.; Tanaka, S. Bacterial Microcompartment Organelles: Protein Shell Structure and Evolution. Annu Rev Biophys 2010, 39, 185–205. 10.1146/annurev.biophys.093008.131418.

(11) McDowell, H. B.; Hoiczyk, E. Bacterial Nanocompartments: Structures, Functions, and Applications. Journal of Bacteriology 2022, 204 (3), e00346–21. 10.1128/jb.00346-21.

(12) Yeates, T. O.; Thompson, M. C.; Bobik, T. A. The Protein Shells of Bacterial Microcompartment Organelles. Curr Opin Struct Biol 2011, 21 (2), 223–231. 10.1016/j.sbi.2011.01.006.

(13) Bobik, T. A.; Havemann, G. D.; Busch, R. J.; Williams, D. S.; Aldrich, H. C. The Propanediol Utilization (Pdu) Operon ofSalmonella Enterica Serovar Typhimurium LT2 Includes Genes Necessary for Formation of Polyhedral Organelles Involved in Coenzyme B12-Dependent 1,2-Propanediol Degradation. Journal of Bacteriology 1999, 181 (19), 5967–5975. 10.1128/jb.181.19.5967-5975.1999.

(14) Cheng, S.; Sinha, S.; Fan, C.; Liu, Y.; Bobik, T. A. Genetic Analysis of the Protein Shell of the Microcompartments Involved in Coenzyme B12-Dependent 1,2-Propanediol Degradation by Salmonella. J Bacteriol 2011, 193 (6), 1385–1392. 10.1128/JB.01473-10.

(15) Liu, T.; Luo, H.; Gao, F. Position Preference of Essential Genes in Prokaryotic Operons. PLOS ONE 2021, 16 (4), e0250380. 10.1371/journal.pone.0250380.

(16) Stewart, K. L.; Stewart, A. M.; Bobik, T. A. Prokaryotic Organelles: Bacterial Microcompartments in E. Coli and Salmonella. EcoSal Plus 9 (1), 10.1128/ecosalplus.ESP-0025-2019. https://doi.org/10.1128/ecosalplus.esp-0025-2019.

(17) Mills, C. E.; Waltmann, C.; Archer, A. G.; Kennedy, N. W.; Abrahamson, C. H.; Jackson, A. D.; Roth, E. W.; Shirman, S.; Jewett, M. C.; Mangan, N. M.; Olvera de la Cruz, M.; TullmanErcek, D. Vertex Protein PduN Tunes Encapsulated Pathway Performance by Dictating Bacterial Metabolosome Morphology. Nat Commun 2022, 13 (1), 3746. 10.1038/s41467-022-31279-3.

(18) Datsenko, K. A.; Wanner, B. L. One-Step Inactivation of Chromosomal Genes in Escherichia Coli K-12 Using PCR Products. Proc Natl Acad Sci U S A 2000, 97 (12), 6640–6645. 10.1073/pnas.120163297.

(19) Sinha, S.; Cheng, S.; Fan, C.; Bobik, T. A. The PduM Protein Is a Structural Component of the Microcompartments Involved in Coenzyme B12-Dependent 1,2-Propanediol Degradation by Salmonella Enterica. J Bacteriol 2012, 194 (8), 1912–1918. 10.1128/JB.06529-11.

(20) Del Duca, S.; Semenzato, G.; Esposito, A.; Liò, P.; Fani, R. The Operon as a Conundrum of Gene Dynamics and Biochemical Constraints: What We Have Learned from Histidine Biosynthesis. Genes 2023, 14 (4), 949. 10.3390/genes14040949.

(21) Schmid, M. B.; Roth, J. R. Gene Location Affects Expression Level in Salmonella Typhimurium. Journal of Bacteriology 1987, 169 (6), 2872–2875. 10.1128/jb.169.6.2872-2875.1987.

(22) Wells, J. N.; Bergendahl, L. T.; Marsh, J. A. Operon Gene Order Is Optimized for Ordered Protein Complex Assembly. Cell Rep 2016, 14 (4), 679–685. 10.1016/j.celrep.2015.12.085.

(23) Ochoa, J. M.; Dershwitz, P.; Schappert, M.; Sinha, S.; Herring, T. I.; Yeates, T. O.; Bobik, T. A. A Single Shell Protein Plays a Major Role in Choline Transport across the Shell of the Choline Utilization Microcompartment of Escherichia Coli 536. Microbiology 2023, 169 (11), 001413. 10.1099/mic.0.001413.

(24) Kast, P. pKSS — A Second-Generation General Purpose Cloning Vector for Efficient Positive Selection of Recombinant Clones. Gene 1994, 138 (1), 109–114. 10.1016/0378-1119(94)90790-0.

(25) Yang, M.; Simpson, D. M.; Wenner, N.; Brownridge, P.; Harman, V. M.; Hinton, J. C. D.; Beynon, R. J.; Liu, L.-N. Decoding the Stoichiometric Composition and Organisation of Bacterial Metabolosomes. Nat Commun 2020, 11 (1), 1976. 10.1038/s41467-020-15888-4.

(26) Kennedy, N. W.; Ikonomova, S. P.; Slininger Lee, M.; Raeder, H. W.; Tullman-Ercek, D. Self-Assembling Shell Proteins PduA and PduJ Have Essential and Redundant Roles in Bacterial Microcompartment Assembly. Journal of Molecular Biology 2020, 433 (2). 10.1016/j.jmb.2020.11.020.

(27) Sawicki, Eugene.; Hauser, T. R.; Stanley, T. W.; Elbert, Walter. The 3-Methyl-2-Benzothiazolone Hydrazone Test. Sensitive New Methods for the Detection, Rapid Estimation, and Determination of Aliphatic Aldehydes. Anal. Chem. 1961, 33 (1), 93–96. 10.1021/ac60169a028.

(28) Yang, M.; Wenner, N.; Dykes, G. F.; Li, Y.; Zhu, X.; Sun, Y.; Huang, F.; Hinton, J. C. D.; Liu, L.-N. Biogenesis of a Bacterial Metabolosome for Propanediol Utilization. Nat Commun 2022, 13 (1), 2920. 10.1038/s41467-022-30608-w.

(29) Cheng, S.; Sinha, S.; Fan, C.; Liu, Y.; Bobik, T. A. Genetic Analysis of the Protein Shell of the Microcompartments Involved in Coenzyme B12-Dependent 1,2-Propanediol Degradation by Salmonella. Journal of Bacteriology 2011, 193 (6), 1385–1392. 10.1128/jb.01473-10.

(30) Kennedy, N. W.; Mills, C. E.; Abrahamson, C. H.; Archer, A. G.; Shirman, S.; Jewett, M. C.; Mangan, N. M.; Tullman-Ercek, D. Linking the Salmonella Enterica 1,2-Propanediol Utilization Bacterial Microcompartment Shell to the Enzymatic Core via the Shell Protein PduB. Journal of Bacteriology 2022, 204 (9), e00576–21. 10.1128/jb.00576-21.

(31) Dank, A.; Zeng, Z.; Boeren, S.; Notebaart, R. A.; Smid, E. J.; Abee, T. Bacterial Microcompartment-Dependent 1,2-Propanediol Utilization of Propionibacterium Freudenreichii. Front Microbiol 2021, 12, 679827. 10.3389/fmicb.2021.679827.

(32) Cheng, C. C.; Duar, R. M.; Lin, X.; Perez-Munoz, M. E.; Tollenaar, S.; Oh, J.-H.; van Pijkeren, J.-P.; Li, F.; van Sinderen, D.; Gänzle, M. G.; Walter, J. Ecological Importance of Cross-Feeding of the Intermediate Metabolite 1,2-Propanediol between Bacterial Gut Symbionts. Appl Environ Microbiol 2020, 86 (11), e00190–20. 10.1128/AEM.00190-20.

(33) Sampson, E. M.; Bobik, T. A. Microcompartments for B12-Dependent 1,2-Propanediol Degradation Provide Protection from DNA and Cellular Damage by a Reactive Metabolic Intermediate. J Bacteriol 2008, 190 (8), 2966–2971. 10.1128/JB.01925-07.

(34) Möker, N.; Brocker, M.; Schaffer, S.; Krämer, R.; Morbach, S.; Bott, M. Deletion of the Genes Encoding the MtrA–MtrB Two-Component System of Corynebacterium Glutamicum Has a Strong Influence on Cell Morphology, Antibiotics Susceptibility and Expression of Genes Involved in Osmoprotection. Molecular Microbiology 2004, 54 (2), 420–438. 10.1111/j.1365-2958.2004.04249.x.

(35) Kennedy, N. W.; Ikonomova, S. P.; Slininger Lee, M.; Raeder, H. W.; Tullman-Ercek, D. Self-Assembling Shell Proteins PduA and PduJ Have Essential and Redundant Roles in Bacterial Microcompartment Assembly. J Mol Biol 2021, 433 (2), 166721. 10.1016/j.jmb.2020.11.020.

(36) Parsons, J. B.; Frank, S.; Bhella, D.; Liang, M.; Prentice, M. B.; Mulvihill, D. P.; Warren, M. J. Synthesis of Empty Bacterial Microcompartments, Directed Organelle Protein Incorporation, and Evidence of Filament-Associated Organelle Movement. Molecular Cell 2010, 38 (2), 305–315. 10.1016/j.molcel.2010.04.008.

(37) Jakobson, C. M.; Tullman-Ercek, D.; Slininger, M. F.; Mangan, N. M. A Systems-Level Model Reveals That 1,2-Propanediol Utilization Microcompartments Enhance Pathway Flux through Intermediate Sequestration. PLoS Comput Biol 2017, 13 (5), e1005525. 10.1371/journal.pcbi.1005525.

(38) Lehman, B. P.; Chowdhury, C.; Bobik, T. A. The N Terminus of the PduB Protein Binds the Protein Shell of the Pdu Microcompartment to Its Enzymatic Core. J Bacteriol 2017, 199 (8), e00785–16. 10.1128/JB.00785-16.

(39) Fan, C.; Bobik, T. A. The N-Terminal Region of the Medium Subunit (PduD) Packages Adenosylcobalamin-Dependent Diol Dehydratase (PduCDE) into the Pdu Microcompartment. J Bacteriol 2011, 193 (20), 5623–5628. 10.1128/JB.05661-11.

(40) Chowdhury, C.; Chun, S.; Sawaya, M. R.; Yeates, T. O.; Bobik, T. A. The Function of the PduJ Microcompartment Shell Protein Is Determined by the Genomic Position of Its Encoding Gene. Molecular Microbiology 2016, 101 (5), 770–783. 10.1111/mmi.13423.

(41) Rillema, R.; Hoang, Y.; MacCready, J. S.; Vecchiarelli, A. G. Carboxysome Mispositioning Alters Growth, Morphology, and Rubisco Level of the Cyanobacterium Synechococcus Elongatus PCC 7942. mBio 2021, 12 (4), 10.1128/mbio.02696-20. https://doi.org/10.1128/mbio.02696-20.

(42) Zaslaver, A.; Mayo, A.; Ronen, M.; Alon, U. Optimal Gene Partition into Operons Correlates with Gene Functional Order. Phys. Biol. 2006, 3 (3), 183. 10.1088/1478-3975/3/3/003.

