## Supplementary File for "Genetic Programming of Bacterial Microcompartments: Operon Order as a Tool for Nanoscale Morphogenesis"

##### Comparison of bacterial microcompartment-encoding operons

Bacterial microcompartments (BMCs) are protein-based organelles found in many bacteria that encapsulate specific metabolic pathways. Unlike membrane-bound organelles, BMCs are made entirely of shell proteins that assemble into a semi-permeable structure. All bacterial microcompartments are composed of combinations of shell proteins, BMC-P (pentamer protein), BMC-H (hexamer protein), BMC-Ts (standard trimer protein) and BMC-Tdp (double pore trimer protein) with their respective enzyme proteins. A comparison of BMC-P protein is done using a webtool called BMC caller, locus database to analyze the position of BMC-P across BMCs **(Table 1)**, and we have observed that BMC-P is interspaced within the shell protein and enzyme protein encoding genes, somewhere in the middle of the operons.

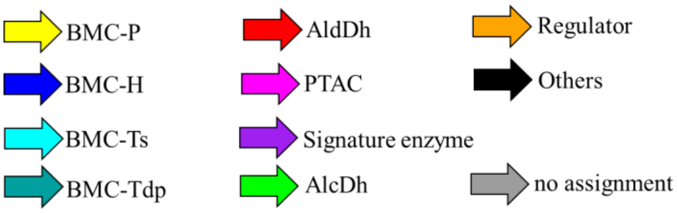


| BMC locus type | Description | Operon structure |
|---|---|---|
| ACI | Acidobacterium microcompartment | 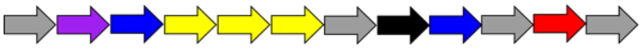 |
| ARO | Aromatic substrate microcompartment | 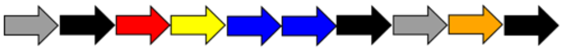 |
| BUF1 | Bacterial microcompartment of unknown function (and no AldDh) | 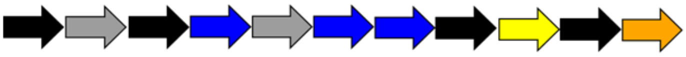 |
| CsomeACh | Alpha-carboxysome of che moautotroph bacteria | 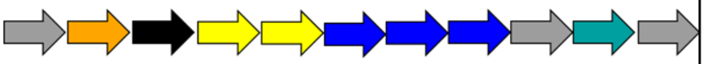 |
| CsomeACy | Alpha-carboxysome of cyan obacteria | 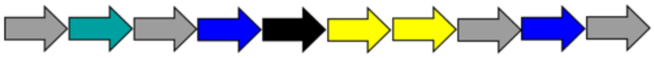 |

| | | |
|---|---|---|
| **CsomeBCy** | **Beta- carboxysome of cyan obacteria** | 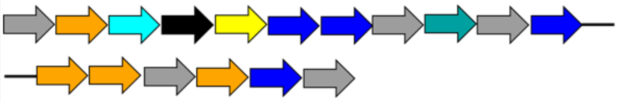 |
| **ETU** | **Ethanol utilizing microcompartment** | 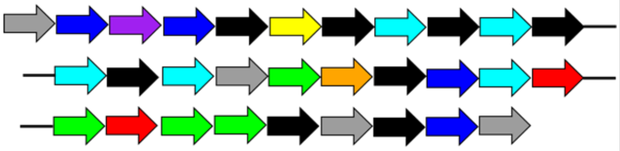 |
| **EUT** | **Ethanolamine utilizin g microcompartment** | 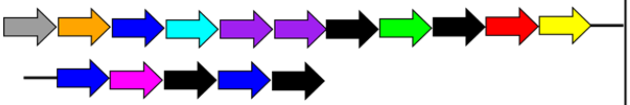 |
| **GRM1A** | **Glycyl radical enzyme containing microcom partment** | 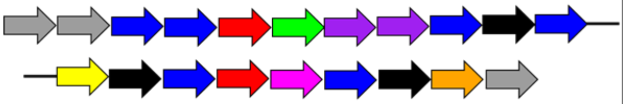 |
| **PDU1AB** | **Propanediol utilizatio n microcompartment** | 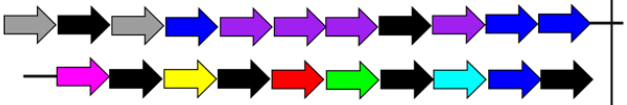 |
| **PVM** | **Planctomycete and Verrucomicrobia microcompartment** | 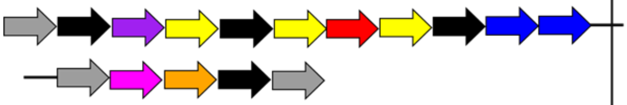 |

**Table 1: List of some of the known Bacterial microcompartment types with their respective operon structures. Operons and notations are adapted using the BMC locus database from the BMC caller webtool[1].**

**Rationale and strategy of *pduN* gene repositioning within the operon**

The 1,2-propanediol utilization bacterial microcompartment (PduBMC) is a protein-based organelle found in *Salmonella enterica* and some other enteric bacteria that facilitates the metabolism of 1,2-propanediol. It is composed of a selectively permeable polyhedral shell formed by multiple shell proteins that encapsulate enzymes involved in the metabolic pathway. This compartmentalization enhances metabolic efficiency while preventing the cytosolic release of toxic intermediates such as propionaldehyde. The assembly of the PduBMC is genetically encoded by the *pdu* operon, which contains genes for shell proteins,

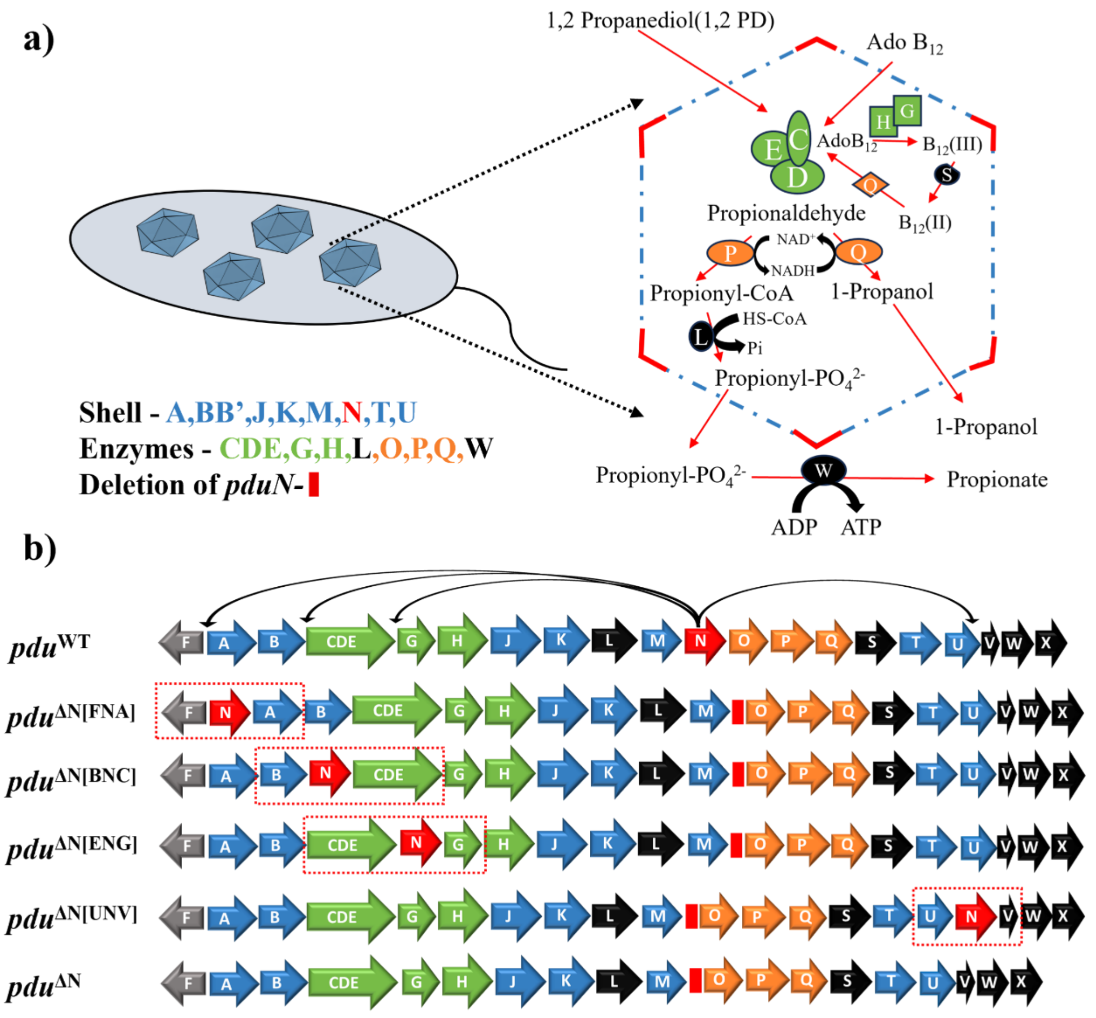


**Supplementary figure 1: Rationale and strategy of *pduN* gene repositioning within the operon. General structure and metabolic function of PduBMC, the outer polyhedral shell is made up of the mentioned shell proteins and encapsulated enzymes systematically metabolize 1,2 propanediol in a B12-dependent manner a) Structure of the *pdu* operon and the marked repositioned mutants generated in this study, upstream of primary shell protein gene *pduA* [*pdu*$^{\Delta N[FNA]}$], preceding adjacent to the primary signature enzyme *pduCDE* [*pdu*$^{\Delta N[BNC]}$], downstream of *pduCDE* [*pdu*$^{\Delta N[ENG]}$], and at the operon terminus following all shell protein genes [*pdu*$^{\Delta N[UNV]}$] b).**

encapsulated enzymes, and accessory factors required for its formation and function **(Supp Figure 1a)**. In *pdu* operon *pduN* occupies a mid-operon position, raising the possibility that its genomic location may influence assembly processes. Based on this observation, we hypothesized that gene position could contribute to structural and functional outcomes during nanoscale morphogenesis. To test this, we treated operon sequence as an engineerable variable and systematically repositioned *pduN* within the *pdu* operon. The following four variants were generated: (i) upstream of primary shell protein gene *pduA* [*pdu*$^{\Delta N[FNA]}$], (ii) preceding adjacent to the primary signature enzyme *pduCDE* [*pdu*$^{\Delta N[BNC]}$], (iii) downstream of *pduCDE* [*pdu*$^{\Delta N[ENG]}$], and (iv) at the operon terminus following all shell protein genes [*pdu*$^{\Delta N[UNV]}$]. All variants were constructed using λ-Red recombination methodology with selection mediated by the *mPhes-gent* cassette. These constructs were introduced into a deleted *pduN* (*pdu*$^{\Delta N}$) strain to isolate the impact of native gene position on BMC morphology, assembly, and metabolic activity. Wild-type (*pdu*$^{WT}$) and *pdu*$^{\Delta N}$ individual strains were included as controls **(Supp Figure 1b)**.

**Agarose gel electrophoresis was performed to confirm the successful repositioning of the *pduN* gene within the operon and its subsequent deletion from the native locus**

The strains generated for this study were confirmed by using PCR method with position specific upstream and downstream primer sets. For example, *pdu*$^{\Delta N[FNA]}$ was confirmed with forward primer which binds to *pduF* that is upstream to *PduA* and reverse primer binds within *pduBB'*. The increase in band size in *pdu*$^{\Delta N[FNA]}$ PCR result compared to *pdu*$^{WT}$ highlight the incorporation of *pduN* upstream to *pduA* in the operon. Similarly, a decrease in band size of all mutants compared to *pdu*$^{WT}$ suggests deletion of *pduN* from its native position **(Supp Figure 2)**.

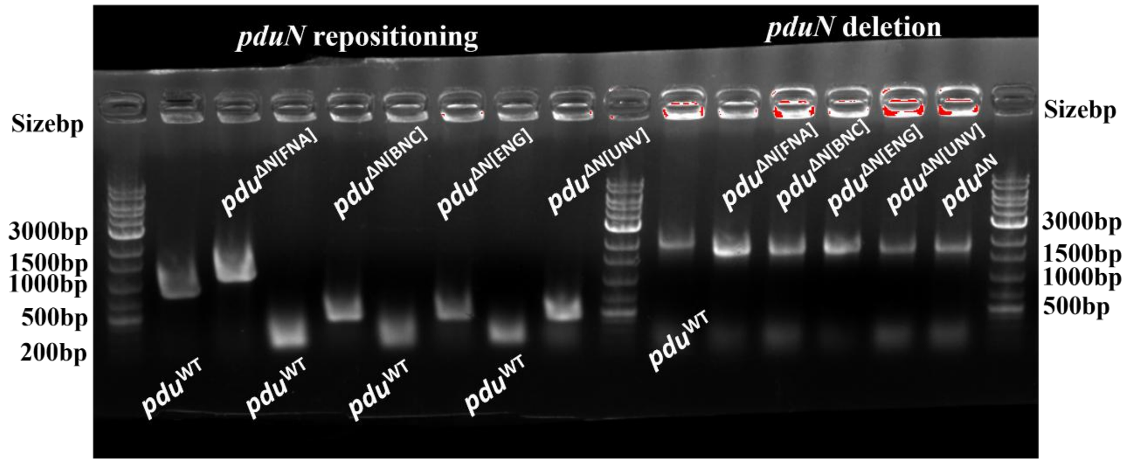


**Supplementary Figure 2: Agarose gel electrophoresis was performed to confirm the successful repositioning of the *pduN* gene within the operon and its subsequent deletion from the native locus. PCR amplification using primers flanking the targeted site revealed a size shift in the bands: an increased band size indicated the insertion of *pduN* at the new position, whereas a reduced band size confirmed its deletion from the original location.**

__Influence of gene order on BMC Expression__

To assess whether repositioning of *pduN* affected BMC biogenesis, we analyzed BMC-induced cell lysates from all engineered strains. Cultures were grown in minimal medium (NCE supplemented with 1,2-propanediol) to induce BMC formation, and lysates were examined by SDS-PAGE for  profiling. All variants displayed protein bands corresponding in molecular weight to the core shell protein PduBB′ and the catalytic enzyme complex PduCDE, indicating that gene repositioning does not disrupt basal protein expression.

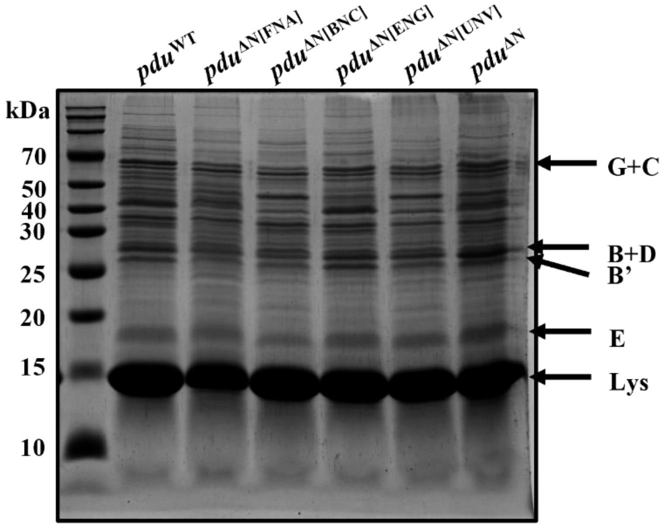


**Supplementary Figure 3: SDS-PAGE gel images of bacterial cell lysates expressing different mutant BMCs, some of the major shell and enzyme protein bands are labelled to check the expression of BMC proteins.**

#### <u>Gene expression study of PduBMC shell proteins</u>

To understand the assembly of BMC, we have done time-dependent RT-qPCR of the first gene in the operon (*pduA*) and the repositioned gene (*pduN*) in WT and mutant BMCs **(Sup Figure 3)**. In all the repositioned mutants we have observed, the expression of both genes, i.e. *pduA* and *pduN,* at different time points. Although the level of expression varies in different mutants with time, such as after 2 hours, *pduN* transcription is enhanced several-fold in *pdu*$^{\Delta N[FNA]}$ mutant. Further, a sharp increase of both *pduA* and *pduN* transcription is found to be present in *pdu*$^{\Delta N[UNV]}$ mutant after 6 hours. Similarly, enhanced expression of *pduN* is observed in *pdu*$^{\Delta N[FNA]}$, *pdu*$^{\Delta N[BNC]}$, and *pdu*$^{\Delta N[ENG]}$ mutants after 14 hours of induction. As the regulation of gene expression is dependent on various factors, including the presence of neighbouring genes, analyses of these changes are difficult. Further, no literature suggests the incorporation of protein expressed from the *pdu* operon is directly proportional to the abundance of proteins in the PduBMC structure. Together, understanding the mechanistic connection between gene order and assembly is crucial and really important to understand PduBMC biogenesis, but it cannot be elucidated from this study.

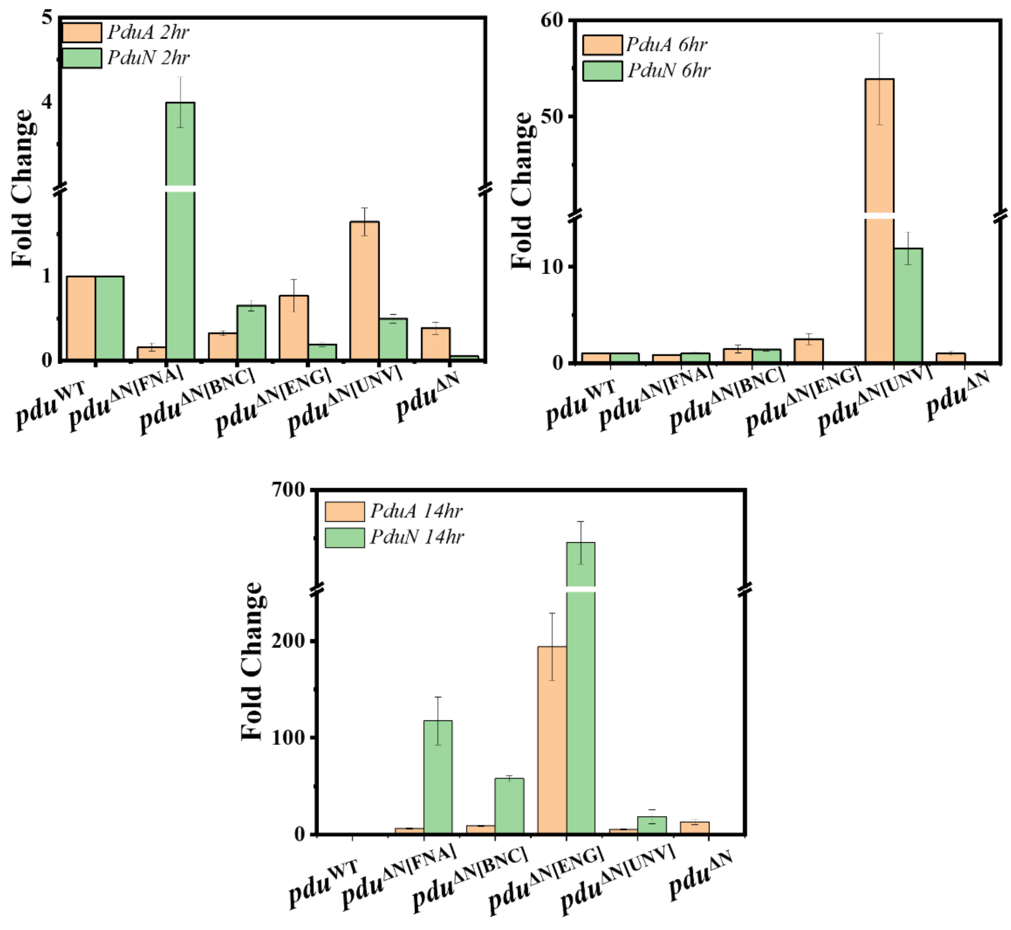


**Supplementary Figure 4: Gene expression study of PduBMC shell proteins. RT-qPCR analysis of *pduA* and *pduN* expression at different time points. After 2 hours, *pduN* transcription is enhanced several-fold in *pdu*$^{\Delta N[FNA]}$ mutant. Further, a sharp increase in both *pduA* and *pduN* transcription is found to be present in *pdu*$^{\Delta N[UNV]}$ mutant after 6 hours. Similarly, enhanced expression of *pduN* is observed in *pdu*$^{\Delta N[FNA]}$, *pdu*$^{\Delta N[BNC]}$, and *pdu*$^{\Delta N[ENG]}$ mutants after 14 hours of induction.**

Comparison of the length of bacteria with the length of purified BMC structures

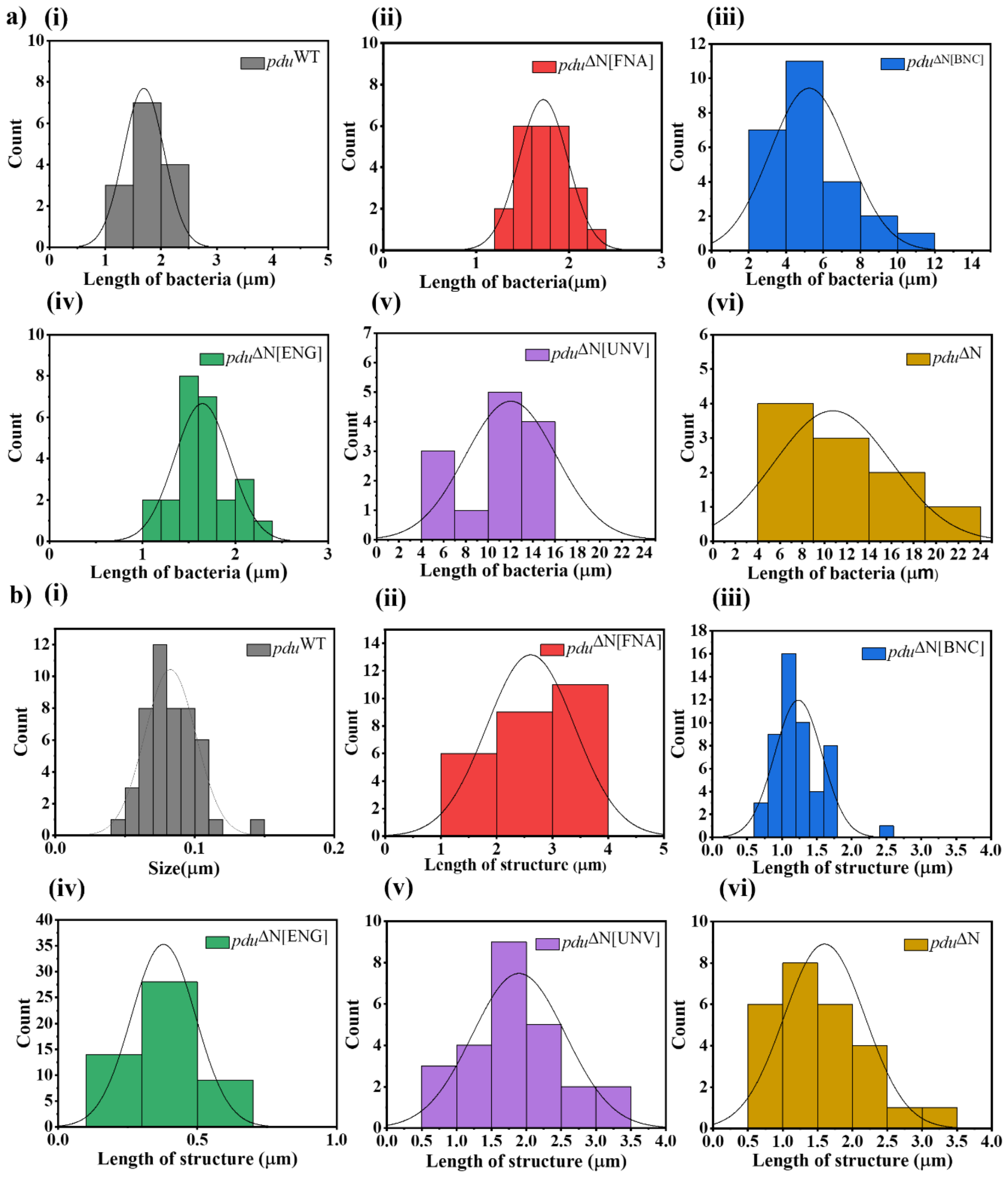


**Supplementary Figure 5: Comparison of length of bacteria with length of purified BMC structures. Length of Bacteria a) and the length of purified structures b) calculated from electron microscopy images using ImageJ, Length of *pdu*$^{WT}$ and *pdu*$^{\Delta N[ENG]}$ structures bi,aiv) were shorter than the size of bacteria ai), whereas *pdu*$^{\Delta N[BNC]}$, *pdu*$^{\Delta N[UNV]}$, and *pdu*$^{\Delta N}$ structures bii,bv, and bvi) had greater or equivalent length that allows these structures to span the entire cell and led to the formation of bacterial chain-like morphology. Moreover, the lengths of BMC mutants expressing bacteria aiii, av, and avi) had a direct correlation with the lengths of structures bii,bv, and bvi).**

The linear alignment of bacteria expressing different mutants in FESEM study **(Figure 7a)** suggests partial or complete inhibition of cell fission, likely due to elongated internal structures traversing the cytoplasm and obstructing septation. To check this, we compared the length of WT bacteria with the purified BMC structure lengths observed by TEM study **(Sup Figure 4).** The length of $pdu^{WT}$ and $pdu^{\Delta N[ENG]}$ structures **bi,aiv)** were observed to be shorter than the size of bacteria **ai)**, whereas $pdu^{\Delta N[BNC]}$, $pdu^{\Delta N[UNV]}$, and $pdu^{\Delta N}$ structures **bii,bv, and bvi)** had greater or equivalent length that allows these structures to span the entire cell and led to the formation of bacterial chain-like morphology.

Further, the lengths of BMC mutants expressing bacteria **aiii, av, and avi)** had a direct correlation with the lengths of structures **bii,bv, and bvi)**. This makes these nanostructures a promising tool to utilize genetically controlled systems for customized applications **(Figure 7b)**.

### <u>List of primers used in the study</u>

| Name | Sequence (5'→3') |
|---|---|
| **FPFmphesB** | gcatcttcttatagtcccaactatcggaacactccatgcgaggtctttcgttattaggtggcggtactt |
| **RPFmphesB** | cacacgggcaatcacctgcgccatgatctgttccaccagctcattgctgctcattggctaattcccctagaagcggccgcgaattc |
| **FP-FNA** | ctcgcatcttcttatagtcccaactatcggaacactccatgcgagg |
| **RP-FNA** | cacacgggcaatcacctgcgccatgatctgttccaccagctcattgctgctcattggctaattcccttcggt |
| **FPBmphesC** | Cttgcgaccctcggttctgaaccgaaaaacgatcgtccgtcctacatctgacgttattaggtggcggtactt |
| **RPBmphesC** | tttcgccagtgcttcaaatcttttcgatctcatgaatcagcctcgtgggtagaattcgcggccgcttctag |
| **FP-BNC** | cttgcgaccctcggttctgaaccgaaaaacgatcgtccgtcctacatctgaatcatgcatctggcacgagtc |
| **RP-BNC** | tttcgccagtgcttcaaatcttttcgatctcatgaatcagcctcgtgggtattaacacgaaagcgtatctac |
| **FPEmphesG** | gaagcggcagggctgtacgttgagcgtaaaaaactcaaaggcgacgattaacgttattaggtggcggtactt |
| **RPEmphesG** | cgttgatgagttaccgatgtcaatgccagctatatatcgcatacgaaatccgaattcgcggccgcttctag |
| **FP-ENG** | gaagcggcagggctgtacgttgagcgtaaaaaactcaaaggcgacgattaaatcatgcatctggcacgagtc |
| **RP-ENG** | cgttgatgagttaccgatgtcaatgccagctatatatcgcatacgaaatccttaacacgaaagcgtatctac |
| **FPUmphesV** | acgctgggcgaaatgatgcagttcaccacttgctcgatcacccggacgtaacgttattaggtggcggtactt |
| **RPUmphesV** | cgaggtttttccgcactggctggggccgataaacatcaaacgcttcatgactgaattcgcggccgcttctag |
| **FP-UNV** | acgctgggcgaaatgatgcagttcaccacttgctcgatcacccggacgtaaatcatgcatctggcacgagtc |
| **RP-UNV** | cgaggtttttccgcactggctggggccgataaacatcaaacgcttcatgactttaacacgaaagcgtatctac |
| **FPΔN** | gcgcgtgacgcggccaatgcgcggaatattcaattaattaagcaggagtaacgttattaggtggcggtactt |
| **RPΔN** | tgatgtggtgccagcgtcacctgttcgggtataaatcgccataaccgccccgaattcgcggccgcttctag |
| **FP-N Oligo** | gcgcgtgacgcggccaatgcgcggaatattcaattaattaagcaggagtaaattgtagatacgctttcgtgttaa |
| **RP-N Oligo** | tgatgtggtgccagcgtcacctgttcgggtataaatcgccataaccgccccttaacacgaaagcgtatctacaat |
| **FP-BNC REV** | gtgaagcggcagggctgtacgttgagcgtaaaaaactcaaaggcgacgattaaggatttc |
| **RP-BNC REV** | cttccgttgatgagttaccgatgtcaatgccagctatatatcgcatacgaaatccttaaT |
| **FP-ENG REV** | aaaaccgtccttgcgaccctcggttctgaaccgaaaaacgatcgtccgtcctacatctga |
| **RP-ENGREV** | cgccagtgcttcaaatcttttcgatctcatgaatcagcctcgtgggtatcagatgtagga |

**Table 2: List of primers used to perform gene repositioning to generate repositioned, deleted and reversed mutants using *mphes-gent* cassette.**

| Name | Sequence (5'→3') |
|---|---|
| **PduDFwd** | aggagatatacatatggcagatctatggaaattaatgaaaaa |
| **Int-DsfGFPF** | gacgtactccgcgatatgaagtccaaaggtgaagaactgttc |
| **Int-DsfGFPR** | gaacagttcttcacctttggacttcatatcgcggagtacgtc |
| **sfGFPRev** | ggtttctttaccagactcgagtcatttgtagagctcatccatgcc |

**Table 3: List of primers used to clone D-sfGFP in the pLac22 plasmid to track BMCs within bacteria in this study.**

| Plasmid Name | Use |
|---|---|
| **pKD46** | To perform λ Red Recombination |
| **pMG1** | To amplify *mphes-gent* cassette |
| **pLac22** | To clone D-sfGFP |
| **pET14-GFP30_Encap** | To amplify sf-GFP |

**Table 4: List of plasmids used in the study with their uses.**

**Strains developed in this study**

| Strain No. | Strain Name | Strain details |
|---|---|---|
| SS1001 | $pdu^{\Delta N[FNA]}$ | *pduN is* placed between the *pduF* (propanediol facilitator) and *pduA* (first shell protein gene) with *pduN* deleted from its original place. |
| SS1002 | $pdu^{\Delta N[BNC]}$ | *pduN* placed in between the *pduBB'* (Trimeric shell protein gene) and *pduCDE* (signature enzyme gene), with *pduN* being deleted from its original place. |
| SS1003 | $pdu^{\Delta N[ENG]}$ | *pduN is* placed between the *pduCDE* (signature enzyme gene) and *pduG* (gene encoding the enzyme that helps in recycling of Ado-$B_{12}$) with *pduN* deleted from its original place. |
| SS1004 | $pdu^{\Delta N[UNV]}$ | *pduN* placed in between the *pduU* (last known shell protein encoding gene) and *pduV* (function not defined) with *pduN* is deleted from its original place. |
| SS1005 | $pdu^{\Delta N}$ | *pduN* is deleted from its original place and replaced by an oligo having native RBS for downstream gene, *pduN* is deleted from its original place. |
| SS1006 | Reverse $pdu^{\Delta N[BNC]}$ | *pduN* is placed to its native position. |
| SS1007 | *Reverse* $pdu^{\Delta N[ENG]}$ | *pduN* is placed to its native position. |

*All the strains were developed in *Salmonella enterica* subsp. *enterica* serovar Typhimurium (LT2).

References

(1) Sutter, M.; Kerfeld, C. A. BMC Caller: A Webtool to Identify and Analyze Bacterial Microcompartment Types in Sequence Data. *Biol Direct* **2022**, *17* (1), 9. https://doi.org/10.1186/s13062-022-00323-z.
